# The regulatory landscape of 5′ UTRs in translational control during zebrafish embryogenesis

**DOI:** 10.1101/2023.11.23.568470

**Authors:** Madalena M. Reimão-Pinto, Sebastian M. Castillo-Hair, Georg Seelig, Alex F. Schier

## Abstract

The 5′ UTRs of mRNAs are critical for translation regulation, but their *in vivo* regulatory features are poorly characterized. Here, we report the regulatory landscape of 5′ UTRs during early zebrafish embryogenesis using a massively parallel reporter assay of 18,154 sequences coupled to polysome profiling. We found that the 5′ UTR is sufficient to confer temporal dynamics to translation initiation, and identified 86 motifs enriched in 5′ UTRs with distinct ribosome recruitment capabilities. A quantitative deep learning model, DaniO5P, revealed a combined role for 5′ UTR length, translation initiation site context, upstream AUGs and sequence motifs on *in vivo* ribosome recruitment. DaniO5P predicts the activities of 5′ UTR isoforms and indicates that modulating 5′ UTR length and motif grammar contributes to translation initiation dynamics. This study provides a first quantitative model of 5′ UTR-based translation regulation in early vertebrate development and lays the foundation for identifying the underlying molecular effectors.

**Highlights:**

- *In vivo* MPRA systematically interrogates the regulatory potential of endogenous 5′ UTRs
- The 5′ UTR alone is sufficient to regulate the dynamics of ribosome recruitment during early embryogenesis
- The MPRA identifies 5′ UTR *cis*-regulatory motifs for translation initiation control
- 5′ UTR length, upstream AUGs and motif grammar contribute to the differential regulatory capability of 5′ UTR switching isoforms

## Introduction

Translation starts with the recruitment of the ribosome to an mRNA. In eukaryotes, it typically requires the attachment of the ribosomal pre-initiation complex (PIC) to the 5′ cap of the transcript, followed by PIC scanning of the 5′ untranslated region (5′ UTR) until it reaches a suitable start codon. Only then is a translationally competent ribosome assembled that can initiate protein synthesis (Merrick & Pavitt, 2018). High rates of initiation result in higher ribosome density along the transcript and increased protein synthesis output (Hershey *et al*, 2019; King & Gerber, 2016). Thus, the 5′ UTR serves as a point of control for selective mRNA translation and translation initiation can be rate-limiting for protein production (Araujo *et al*, 2012; Hinnebusch, 2011; Hinnebusch *et al*, 2016).

Translation is remarkably dynamic during cellular differentiation, where fine-tuning of protein synthesis coordinates self-renewal and cell fate acquisition (Buszczak *et al*, 2014; Saba *et al*, 2021; Teixeira & Lehmann, 2019). Physiological cues can reprogram the availability of canonical initiation factors to promote selective mRNA translation (Proud, 2019). In addition, 5′ UTRs harbor structural elements that impact scanning and mRNA translation efficiency. For example, RNA secondary structure and short upstream open reading frames (uORFs) generally attenuate translation of the main ORF (Hinnebusch *et al*., 2016; Leppek *et al*, 2018; Renz *et al*, 2020). Short sequence motifs in the 5′ UTR can serve as a platform for recruiting regulatory *trans*-factors such as RNA-binding proteins (RBPs) (Corley *et al*, 2020; Hentze *et al*, 2018) and provide a means to rapidly modulate translation.

Additional regulatory complexity arises from isoforms with different 5′ UTRs but the same protein coding sequence. For example, in yeast, 5′ UTR variants arise from alternative transcription start site (TSS) utilization and regulate the timed translation of proteins required for meiosis progression (Cheng *et al*, 2018). The zebrafish maternal-to-zygotic transition (MZT) is also characterized by changes in 5′ UTR isoform expression (Haberle *et al*, 2014; Nepal *et al*, 2013). For about 10% of the promoters expressed during the MZT, maternally deposited mRNAs are initiated from a different position than the zygotically re-expressed counterpart. To what extent such 5′ UTR isoform switching impacts the dynamics of translation is unknown.

More generally, we lack a systematic understanding of the regulatory roles of 5′ UTRs at transcriptomic scale during vertebrate embryogenesis. This in part because translation is influenced by several features of the mRNA, such as codon sequence composition, regulatory elements in the 3′ UTR or modulation of the poly(A) tail length (Despic & Neugebauer, 2018; Teixeira & Lehmann, 2019). Previous studies found that transcripts with 5′ and 3′ UTR variants can display distinct translatability, but could not determine the regulatory contributions of the 5′ UTR sequence alone (Arribere & Gilbert, 2013; Blair *et al*, 2017; Cheng *et al*., 2018; Dieudonne *et al*, 2015; Floor & Doudna, 2016; Sterne-Weiler *et al*, 2013; Tresenrider *et al*, 2021; Wang *et al*, 2016; Wong *et al*, 2016). One approach to comprehensively define the regulatory capability of 5′ UTRs are massively parallel reporter assays (MPRAs) (Castillo-Hair *et al*, 2023; Cuperus *et al*, 2017; Dvir *et al*, 2013; Jia *et al*, 2020; Lim *et al*, 2021; May *et al*, 2023; Niederer *et al*, 2022; Sample *et al*, 2019; Strayer *et al*, 2023). MPRAs allow to interrogate the regulatory output of thousands of pre-defined sequences, independently of other transcript features. For instance, recent MPRAs have identified 3′ UTR sequences that regulate mRNA degradation and polyadenylation during the zebrafish MZT (Rabani *et al*, 2017; Vejnar *et al*, 2019; Xiang *et al*, 2023).

To systematically determine the contribution of zebrafish 5′ UTRs to translation initiation during early embryogenesis, we developed an *in vivo* 5′ UTR MPRA coupled to polysome profiling. Polysome profiling separates mRNAs along a density gradient based on the number of ribosomes bound, which reflects how efficiently they are translated (Cammas *et al*, 2022; Floor & Doudna, 2016). By combining polysome profiling with high-throughput sequencing, we determined the translational dynamics of mRNAs harboring endogenous 5′ UTRs representing over 11,400 transcripts expressed during embryogenesis, including 5′ UTR isoforms arising from alternative TSS usage. Our work shows that the 5′ UTR sequence alone is sufficient to regulate ribosome recruitment dynamics by over a 100-fold range during embryogenesis. The *in vivo* MPRA captures the broad inhibitory impact of 5′ UTR secondary structures and uORFs and defines the translation initiation site consensus context sequence (AAAC<u>AUG</u>) as optimal for translation initiation in zebrafish. Furthermore, it reveals motifs enriched in 5’ UTRs that display distinct translational behaviors during embryogenesis. Dozens of motifs are novel, while others match consensus motifs that are bound by RNA-binding proteins in other species. To quantify the contribution of 5′ UTR features to ribosome recruitment, we used the MPRA data to train a deep learning model, DaniO5P. Our analyses reveal a previously unappreciated overarching effect of 5′ UTR length on ribosome recruitment capability *in vivo*. Finally, we show that 5′ UTR isoforms arising from alternative TSS switching during the zebrafish MZT result in shifts of 5′ UTR length and motif composition that alter their regulatory capabilities. By integrating the contribution of sequence features and 5′ UTR length, DaniO5P predicts the differential regulatory activities of 5’ UTR isoforms.

## Results

### Annotation of 5′ UTRs during zebrafish embryogenesis

The study of 5′ UTR activity for a given gene rests on the precise annotation of TSSs and start codons. However, genome-wide transcriptome analyses do not precisely annotate TSSs, and annotation databases provide reviewed 5′ UTR sequences for only a few thousand zebrafish transcripts (Akirtava *et al*, 2022; Baranasic *et al*, 2022; Lawson *et al*, 2020; Leppek *et al*., 2018). To precisely recover 5′ UTR sequences of genes expressed throughout early embryogenesis, we used a publicly available cap analysis of gene expression (CAGE) dataset encompassing the first 33 hours of zebrafish development (Nepal *et al*., 2013). CAGE determines TSSs of transcribed and 5′ capped mRNAs at single nucleotide resolution (Shiraki *et al*, 2003). We re-mapped zebrafish CAGE data of 12 zebrafish developmental stages (from the unfertilized egg to the prim20 stage) (Nepal *et al*., 2013) to the most recent zebrafish genome annotation (GRCz11). Then, we matched CAGE-recovered TSS genomic coordinates with coding sequence start site coordinates of annotated transcripts to computationally extract 5′ UTR sequences (Figures 1A and S1A; STAR Methods). After filtering steps (see STAR Methods), our computational analyses recovered 13,309 5′ UTR sequences of 10,354 genes expressed during the first 33 hours of zebrafish development (Table S1). The 5′ UTR sequences range from 15 to 2,813 nucleotides (nts) and have a median length of 141 nts (Figure 1B), comparable to the length of previously confidently annotated zebrafish 5′ UTRs (3,904 5′ UTRs with median length of 131 nts) (Leppek *et al*., 2018).

**Figure 1.**
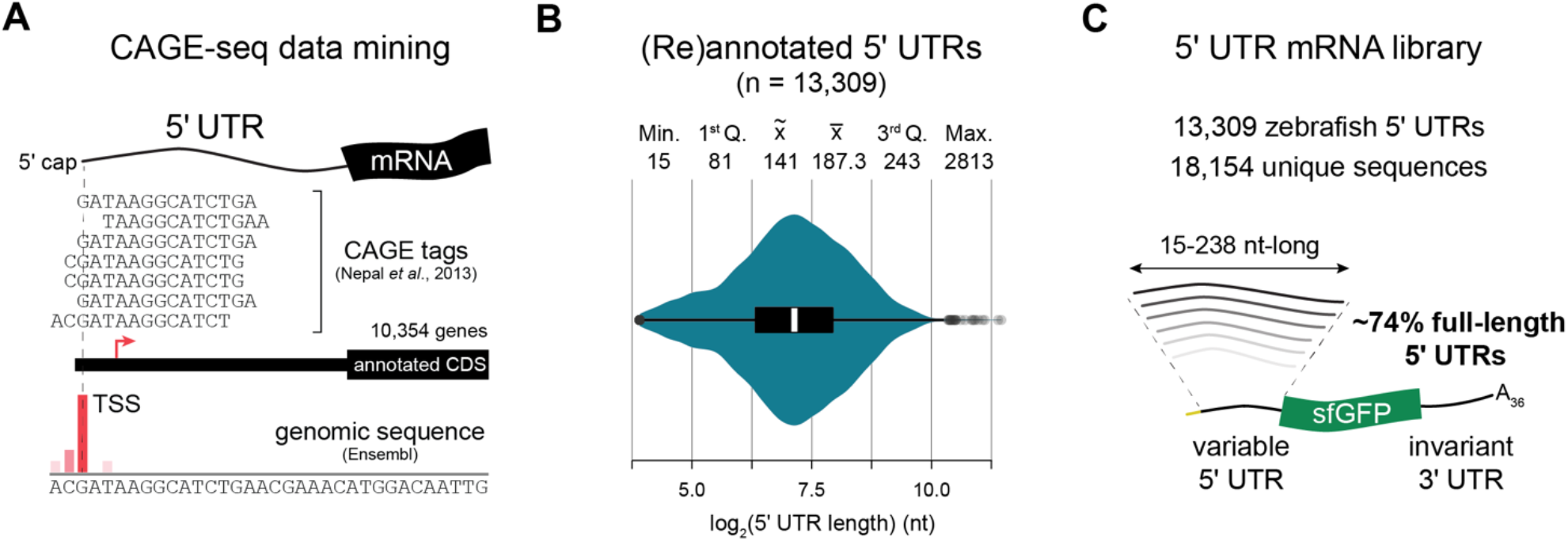
A reporter library for 5’UTR *cis*-regulatory features that regulate translation during zebrafish embryogenesis. (A) Strategy for recovering 5’ UTR sequences. Publicly available cap analysis of gene expression (CAGE) data (Nepal *et al*., 2013) was used to recover transcription start sites (TSSs) and integrated with site genomic positions of annotated coding sequences start sites (Ensembl). (B) Violin plot of the length distribution of 13,309 unique 5’ UTRs represented in the massively parallel reporter assay (MPRA). Center line of boxplot represents median, box limits represent the upper and lower quartiles. Summary statistics of 5’ UTR lengths (in nucleotides) are provided. (C) *In vitro* synthesized 5’ UTR mRNA library with variable 5’ UTR sequence ranging from 15 to 238 nts in length and an invariant sfGFP coding sequence, 3’ UTR and a 36 nt-long poly(A)-tail; of the 18,154 sequences assayed, ∼74% of the 5’ UTRs (n = 9,863) consist of the endogenous uninterrupted 5’ UTR sequence. See also Figure S1 and Table S1.

To recover 5′ UTR isoforms arising from TSS switching during the maternal-to-zygotic transition (Figure S1B), we used CAGEr (Haberle *et al*, 2015). CAGEr determines whether a promoter is significantly shifted either upstream or downstream of the dominant TSS by comparing CAGE signal along individual promoters in two different stages of development (Haberle *et al*., 2015; Haberle *et al*., 2014; Nepal *et al*., 2013). We defined a 5′ UTR switch when at least 60% of the transcription in one stage was independent and happening outside of the region used to initiate transcription from the same promoter in the other developmental stage (shifting score ≥ 0.6, p-value ≤ 0.01, K-S test FDR ≤ 0.01) (STAR Methods). Using this criterium, about 10% of the expressed genes (n = 1,075) scored as 5′ UTR isoform switching (Table S1), with the majority of the switching events occurring during the major wave of zygotic genome activation (ZGA), as previously described (Haberle *et al*., 2014).

### Construction of a 5**′** UTR reporter library

Next, we designed a reporter library using our reannotated 5′ UTRs. We sought to represent the full diversity of endogenous 5′ UTRs, including length differences, and therefore capture potential regulatory sequences in their endogenous sequence context. Thus, 5′ UTRs ranging from 15 to 238 nt (n = 9,863, ∼74% of the sequences assayed) were included as the native endogenous sequence. The remaining longer 5′ UTRs (n = 3,446) were split into shorter sequences to accommodate oligo synthesis limitations (Figure S1C). For transcripts with more than one dominant TSS within a promoter region but that did not score as switching, we considered the longest 5′ UTR isoform (n = 11,620) (Figure S1A). For transcripts undergoing switches in TSS usage, we included all switching unique 5′ UTR variants (n = 2,721) (Figure S1B). This approach resulted in an *in silico* library consisting of 18,154 unique oligonucleotide sequences representing 13,309 unique zebrafish 5′ UTRs (Figure S1F and Table S1).

We synthesized our library as an oligo pool and cloned it into a plasmid, downstream of a SP6 promoter and a common adaptor, and upstream of a superfolder GFP (sfGFP) reporter and a 3′ UTR devoid of known regulatory elements and associated with high mRNA stability (Rabani *et al*., 2017) (Figures S1D and E; STAR Methods). We then generated a translationally competent 5′ UTR mRNA library (Figures 1C), that was transcriptionally capped and included a 36-nt-long poly(A) tail characteristic of adenylated maternal mRNAs (Subtelny *et al*, 2014) (STAR Methods). Our finally library contained almost all of the designed 5′ UTR sequences (sequence representation > 99%, Figure S1F-H). Because each 5′ UTR drives translation of the same reporter transcript, the mRNA library allows to ask how each 5′ UTR impacts translation initiation independently of codon optimality and of regulatory elements in the 3′ UTR.

### *In vivo* zebrafish 5’UTR MPRA uncovers translation initiation regulation

To systematically determine the impact of 5′ UTRs on translation initiation during early zebrafish embryogenesis, we devised an MPRA of zebrafish 5′ UTR sequences coupled to polysome profiling and next-generation sequencing (Figure 2A). We injected the 5′ UTR mRNA library into one-cell stage zebrafish embryos and collected embryos at the 64-cell stage (2 hours post-fertilization, hpf) of the cleavage period, the sphere stage (4 hpf) of the blastula period and the shield (6 hpf) and bud (10 hpf) stages of gastrulation. The reporter library was readily translated, indicated by sfGFP fluorescence already detectable at the 64-cell stage that increased gradually during the time-course (Figure S2A). A transcript’s coding sequence is generally bound by multiple ribosomes simultaneously (polysomes) when efficiently translated (Cammas *et al*., 2022; King & Gerber, 2016). Therefore, to quantify translation of each library reporter, we collected library-injected staged embryos and performed polysome profiling (Figure 2B) (Chasse *et al*, 2017) (STAR Methods). As previously reported (Leesch *et al*, 2023), we observe an increase in polysomes as the embryo develops (Figure 2B). We collected polysome fractions corresponding to monosomes (80S), low molecular weight (LMW) and high molecular weight (HMW) polysomes, as well as total embryo RNA (total) at each time-point (2, 4, 6 and 10 hpf), in triplicates (Figures 2B and S2B). We included the 80S fraction for analysis as it has been shown that monosomes actively contribute to translation (Arava *et al*, 2003; Biever *et al*, 2020; Heyer & Moore, 2016; Schieweck *et al*, 2023; Schneider *et al*, 2022). We then extracted total RNA from each fraction and prepared next-generation sequencing libraries (Figure S2C; STAR Methods). Three time-course experiments were sequenced at large scale (Pearson’s r = 0.75-0.99 between replicates; Figure S2D) and we calculated relative abundances (transcript per million, TPM) for input (total) and polysome fractionated (80S, LMW and HMW) samples throughout the time-course (Figure 2C; Table S1). To determine differential ribosome occupancy of reporter mRNAs, we calculated the ribosome recruitment score (RRS) (Niederer *et al*., 2022) for each 5′ UTR mRNA reporter. The RRS is defined as the ratio between the relative abundance of the reporter in the ribosome-bound fraction (TPM ribosome-bound) and its relative abundance in the input pool (TPM total) (Figure 2C). This calculation yielded 5′ UTR mRNA reporter RRS values for each fraction at each time-point (RRS_80S,_ RRS_LMW_ and RRS_HMW_ at 2, 4, 6 and 10 hpf) (Table S2). The RRS reflects the proportion of reporter transcripts in the mRNA pool that are engaged in active translation. Since reporter mRNAs only differ at the level of the 5′ UTR, higher RRS results from an increased frequency in translation initiation due to regulation by the 5′ UTR sequence. Because RRS is normalized to total reporter abundance at each time-point, possible transcript-specific differences in mRNA stability or library preparation bias for different reporter lengths are factored out.

**Figure 2.**
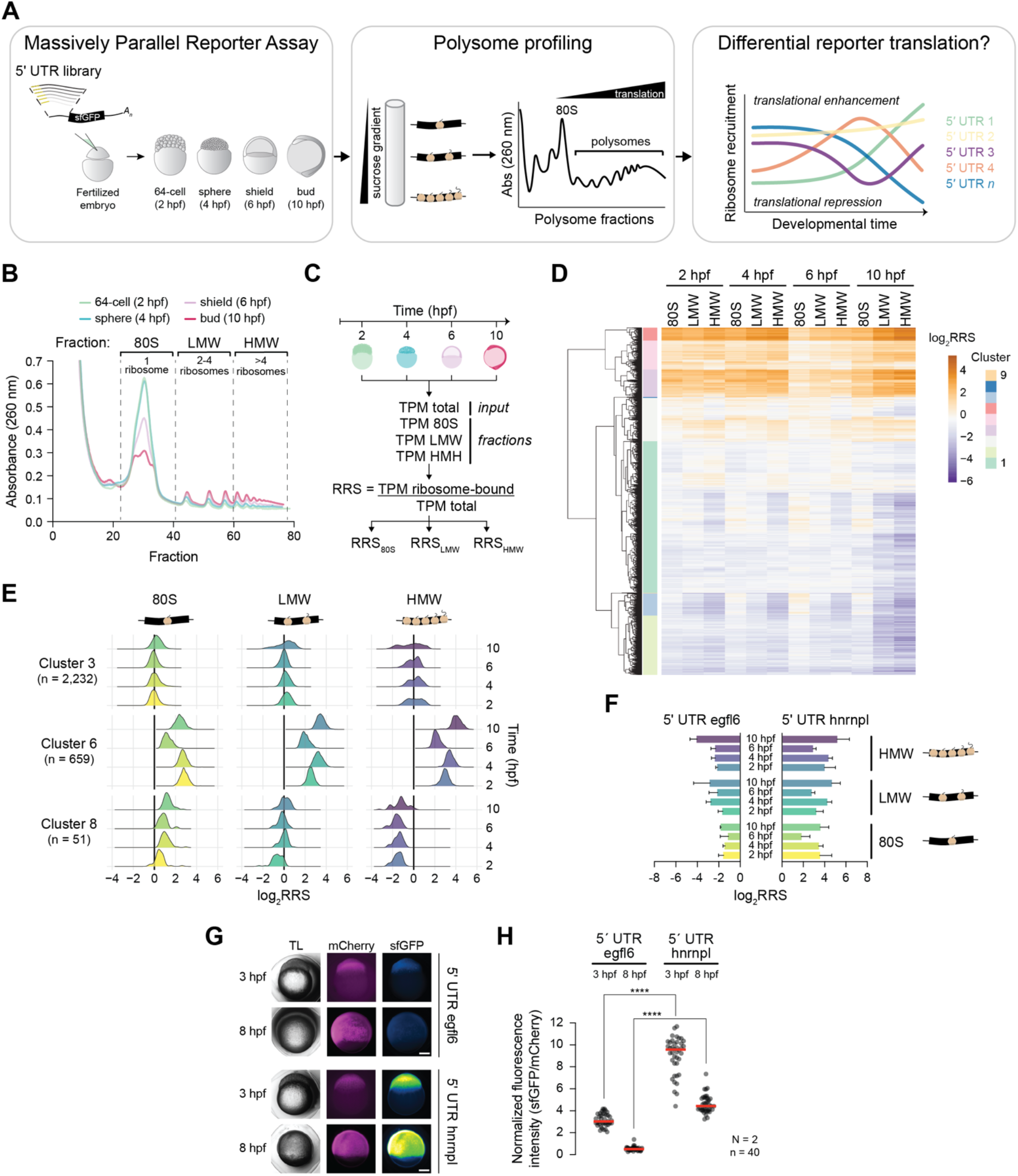
*In vivo* zebrafish 5’UTR MPRA uncovers translation initiation regulation. (A) Schematics of the 5’ UTR massively parallel reporter assay (MPRA) employed. The pooled mRNA reporter library was injected into 1-cell stage embryos and 100 embryos were collected at 2, 4, 6 and 10 hours post-fertilization (hpf). Reporter transcripts were separated based on the number of ribosomes bound by sucrose density gradient centrifugation followed by polysome fractionation. Translational behaviours for each reporter 5’ UTR sequence were determined by high-throughput sequencing. (B) Polysome profiling traces for one of the three replicate experiments. The 80S, low molecular weight (LMW) and high molecular weight (HMW) fractions were collected at 2, 4, 6 and 10 hpf and (C) total RNA was extracted for each of the fractions as well as input (total) sample for quantification of relative 5’ UTR reporter abundance (TPM, transcript per million). Ribosome recruitment scores (RRSs) were calculated for each fraction at each developmental time-point. (D) Heatmap representing mean log2 transformed RRS values of three replicate experiments (n = 17,879). Nine clusters were generated using hierarchical clustering by 5’ UTR sequence (rows) and ordered by fraction and developmental time-point (columns). (E) Density plots of mean log2 transformed RRS values for three representative clusters displaying distinct ribosome recruitment dynamics. (F) Bar plots of mean log2 transformed RRS values for a repressive (egfl6) and an enhancing (hnrnpl) 5’ UTR. Error bars represent SD of three replicate experiments. (G) Fluorescence microscopy images of representative embryos at 3 and 8 hpf that were injected with either the egfl6-5’UTR-sfGFP or the hnrnpl-5’UTR-sfGFP reporter and a control mCherry reporter at the 1-cell stage. TL – transmitted light. (H) Swarm plot displaying relative fluorescence intensities of 3 and 8 hpf embryos injected with each reporter at the 1-cell stage. sfGFP fluorescence was normalized to mCherry intensity in each embryo. Red bars represent median value, each dot represents one embryo (n = 40) from 2 independent experiments (N = 2), **** p-value < 0.0001 Wilcoxon matched-pairs signed rank test. See also Figure S2, Table S2 and STAR Methods.

We observe that the 5′ UTR sequence alone is sufficient to modulate translation initiation during early zebrafish embryogenesis (Figure 2D and S2E). Clustering of RRS values across development shows the translational effects mediated by the 5′ UTR, which can impair (log_2_ RRS < 0) or promote (log_2_ RRS > 0) ribosome recruitment. Importantly, differences in ribosome recruitment are not a mere consequence of differential reporter availability for translation in the mRNA pool (Figure S2F), indicating that active translational regulatory mechanisms are at play. We identify groups of 5′ UTRs that confer different translational behaviors to the reporter mRNA during early embryogenesis (Figure 2E; Table S2). For instance, sequences in Cluster 3 (n = 2,232) lead to relatively constant reporter representation across fractions until 6 hpf, whereas Cluster 6 (n = 659) is composed of 5′ UTRs that promote ribosome recruitment while selectively dampening initiation at the shield (6 hpf) stage. 5′ UTRs in Cluster 8 (n = 51) lead to preferential monosome occupancy of the reporter transcript. These data uncover the regulatory potential that lies in 5′ UTRs and their dynamic regulation throughout embryogenesis.

To test whether RRS is in fact a measure of protein expression, we co-injected either a poorly (log_2_ RRS < 0 for all fractions and time-points; 5′ UTR egfl6) or well (log_2_ RRS > 0; 5′ UTR hnrnlp) ribosome-recruited 5′ UTR-sfGFP reporter with a control mCherry-encoding mRNA and measured relative fluorescence intensities at 3 and 8 hpf (Figures 2F-H). As expected, embryos injected with the 5′ UTR-egfl6-sfGFP-reporter showed a decrease in relative sfGFP intensity as the embryo developed. In contrast, the 5′ UTR-hnrnlp-sfGFP-reporter resulted in significantly higher normalized sfGFP fluorescence intensity at the two time-points (p-value < 0.001, Wilcoxon matched-pairs rank test). These measurements indicate that the rate of reporter translation initiation imposed by the 5′ UTR dictates translational output in this biological context and are in agreement with initiation being a rate-limiting step in protein synthesis. Taken together, the MPRA systematically determined the impact of 5′ UTRs on translation initiation during early zebrafish embryogenesis.

### 5’ UTR regulatory elements shape translation initiation in early development

To analyze our MPRA measurements, we first evaluated the relationship between known 5′ UTR *cis*-acting elements (Hinnebusch, 2011; Hinnebusch *et al*., 2016; Leppek *et al*., 2018) and RRS. Stable structures in the 5′ UTR sequence inhibit scanning and have been shown to impact translation *in vitro* and *in vivo* (Babendure *et al*, 2006; Baim & Sherman, 1988; Cuperus *et al*., 2017; Kozak, 1986a, 1989; Niederer *et al*., 2022; Pelletier & Sonenberg, 1985; Ringner & Krogh, 2005; Wang *et al*, 2022). To determine the relationship between 5′ UTR RNA structure and ribosome occupancy, we predicted 5′ UTR RNA secondary structures using the MXfold2 algorithm (Sato *et al*, 2021). MXfold2 integrates thermodynamic stability and free energy parameters to determine RNA folding scores of a defined sequence using deep learning. We focused our analysis on RRS scores for the HMW fraction (RRS_HMW_) at the bud stage (10 hpf), when global translational activity is high (Leesch *et al*., 2023) (Figure 2B). Plotting RRS scores and MXfold2 RNA folding scores shows that 5′ UTRs with high propensity to form RNA secondary structures are poorly initiated *in vivo* (Figure 3A; Adjusted R^2^ = 0.75, p-value < 2×10^−16^). RRS_HMW_ decreases in a linear fashion as RNA folding score increases for less structured sequences (RNA folding score < 20), whereas more structured 5′ UTRs show no clear relationship with RRS. This suggests a saturation effect of RNA structure on ribosome recruitment. Of note, we observe only a weak correlation between 5′ UTR GC content and RRS_HMW_ (Figure S3A; Adjusted R^2^ = 0.008; p-value < 2×10^−16^), indicating that GC-richness per se does not greatly affect the initiation process *in vivo*.

**Figure 3.**
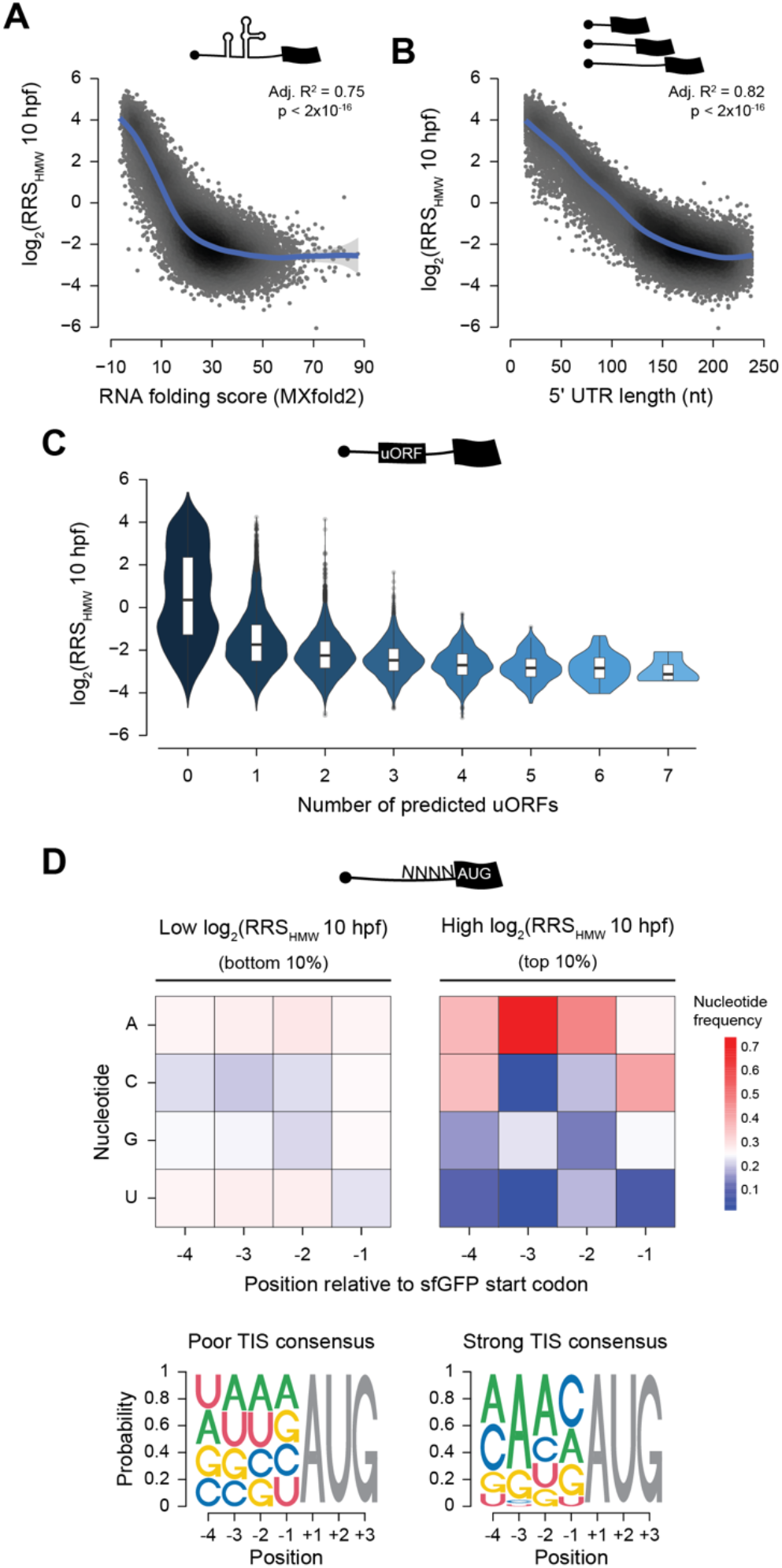
5’ UTR regulatory elements shape translation initiation in early development. (A) Effect of 5’ UTR RNA structures on ribosome recruitment. Scatter plot of RNA folding scores calculated by MXfold2 (x axis) and mean log2 transformed RRS values (y axis) for the HMW fraction at 10 hpf. (B) Effect of 5’ UTR length on ribosome recruitment. Scatter plot of 5’ UTR length in nucleotides (x axis) and mean log2 transformed RRS values (y axis) for the HMW fraction at 10 hpf. Each dot represents one 5’ UTR (n = 17,906) and adjusted R^2^ values for best-fit to a nonlinear generalized additive model and corresponding p-value are presented. (C) Effect of upstream open reading frame (uORF) number on ribosome recruitment. Violin plots of mean log2 transformed RRS values for the HMW fraction at 10 hpf for 5’ UTRs binned by number of predicted uORFs. Center line of boxplots represent median, box limits represent the upper and lower quartiles. (D) Mean nucleotide frequency at positions −4 to - 1 of the reporter canonical sfGFP translation start site for 5’ UTRs with lowest RRS values (bottom 10%, n = 1,791; left panel) and highest RRS values (top 10%, n = 1,791; right panel) for the HMW fraction at 10 hpf. Position weight matrixes of poor and strong TIS consensus context derived from mean nucleotide frequencies are presented. Grey AUG represents the translation start site of sfGFP. See STAR Methods, Figure S3 and Table S3.

The length of the 5′ UTR has been implicated in translation efficiency in mammalian cells, with contradictory observations made between *in vivo* and *in vitro* systems (Bohlen *et al*, 2020; Chappell *et al*, 2006; Kozak, 1988, 1991). To determine the effect of 5′ UTR length on translation *in vivo*, we plotted RRS_HMW_ against 5′ UTR length (Figure 3B). This analysis shows a clear negative relationship between 5′ UTR length and ribosome recruitment in zebrafish embryos (Adjusted R^2^ = 0.82, p-value < 2×10^−16^), revealing a previously unappreciated overarching impact of 5′ UTR length on translation initiation during embryogenesis. Of note, 5′ UTR length and RNA folding score are interdependent and increase in parallel (Figure S3B; Adjusted R^2^ = 0.76, p-value < 2×10^−16^).

Previous transcriptome-wide studies have identified uORFs as prevalent features in vertebrate 5′ UTRs that regulate translation (Calvo *et al*, 2009; Chew *et al*, 2016; Giess *et al*, 2020; Johnstone *et al*, 2016). To evaluate the regulatory impact of 5′ UTR uORFs on ribosome recruitment, we predicted AUG-initiated uORFs using ORFik (Tjeldnes *et al*, 2021). Of the 5′ UTR sequences assayed in the MPRA, around 61% contained predicted AUG-initiated uORFs (STAR Methods; Table S2). We find that 5′ UTRs with a single uORF show significantly lower RRS_HMW_ (p-value < 2.2×10^−16^, Wilcoxon rank sum test) than 5′ UTRs without predicted uORFs (Figure 3C). The presence of additional uORFs further impairs ribosome recruitment in a uORF number dependent manner, albeit to a lesser extent. This result agrees with a broad inhibitory effect of uORFs on the translation of downstream main ORFs, presumably by the detachment of the scanning PIC (Giess *et al*., 2020). Nonetheless, uORF-containing 5′ UTRs display a broad range of RRS_HMW_ values, consistent with a context-dependent effect of uORFs on scanning (Chew *et al*., 2016; Giess *et al*., 2020) and with a combinatorial impact of other 5′ UTR regulatory features on translation initiation. The length-dependency of ribosome recruitment is maintained for 5′ UTRs without predicted uORFs (Figure S3C; Adjusted R^2^ = 0.84, p-value < 2×10^−16^), indicating that the observed length dependency (Figure 3B) cannot be explained by the increased frequency of inhibitory uORFs in longer 5′ UTRs.

To start testing the regulatory importance of sequence elements at large scale, we first focused on the sequence context preceding the translation initiation site (TIS) of the main ORF. Landmark studies identified that this sequence is not random, and that CRCC<u>AUG</u>G (where R denotes a purine) is the optimal consensus sequence for translation initiation (the Kozak sequence) in mammals (Kozak, 1986b, 1987). The zebrafish genome is characterized by the TIS consensus context NRNC<u>AUG</u>G (Grzegorski *et al*, 2014; Hernandez *et al*, 2019; Nakagawa *et al*, 2008). In particular, the sequence context AAAC<u>AUG</u> is associated with high translation rates (Giess *et al*., 2020; Grzegorski *et al*., 2014). To analyze at large scale the effect of the sequence context upstream of the sfGFP main ORF on RRS, we selected reporters from the top 10% and bottom 10% RRS_HMW_ quantiles (Table S3). Then, we determined the identity of the four nucleotides preceding the sfGFP ORF and calculated nucleotide frequencies at each position. Strikingly, this analysis revealed an enrichment of 5′ UTRs with the AAAC<u>AUG</u> sequence context among the top 10% ribosome recruited-reporters (Figure 3D, top 10%). In contrast, the bottom quantile displayed no clear sequence specificity (nucleotide frequency ranging from approximately 20% to 28%), but an increase frequency of uridine (U) at positions −4 to −1 (Figure 3D, bottom 10%) relative to that of all 5′ UTRs assayed (Figure S3D). This observation indicates that the presence of a U at positions −4, −3 or −2 relative to the main ORF is particulary detrimental for translation initiation in zebrafish. Collectively, the MPRA comprehensively quantified the impact of 5′ UTR secondary structure, length, uORFs and TIS sequence context on translation initiation, suggesting that it provides a framework to uncover novel *cis*-acting regulatory elements.

### The *in vivo* MPRA identifies 5’ UTR *cis*-regulatory motifs

The MPRA data (Figure 2) suggests that 5′ UTRs regulate ribosome recruitment dynamically, with amplitude changing over developmental time. We hypothesized that endogenous 5′ UTR sequences carry short sequence motifs that account for these 5′ UTR properties by enhancing or inhibiting ribosome recruitment. To identify such motifs, we took a two-step approach. First, we aimed at identifying short sequence motifs associated with constitutive enhancement or repression of RRS during the developmental time-course (Figure 4A). We performed motif enrichment analysis on sequences that are commonly found in the top (higher 10% RRS) or bottom (lower 10% RRS) quantiles of each of the fractions (80S, LMW and HMW) during the time-course (Figure 4A and S4A; Table S3). For sequence enrichment analysis we used the MEME tool suite (Bailey *et al*, 2015), which enables *de novo* motif discovery and motif scanning of known RBP-bound sequences (Ray *et al*, 2013) (see STAR Methods for search parameters). This analysis yielded 61 unique short-sequence motifs (5-12 nt-long), 30 of which are *de novo* predicted motifs (Table S4) and the remaining match the consensus binding sequence of known RBPs in other species (Dominguez *et al*, 2018; Gerstberger *et al*, 2014; Lambert *et al*, 2014; Ray *et al*., 2013; Van Nostrand *et al*, 2020).

**Figure 4.**
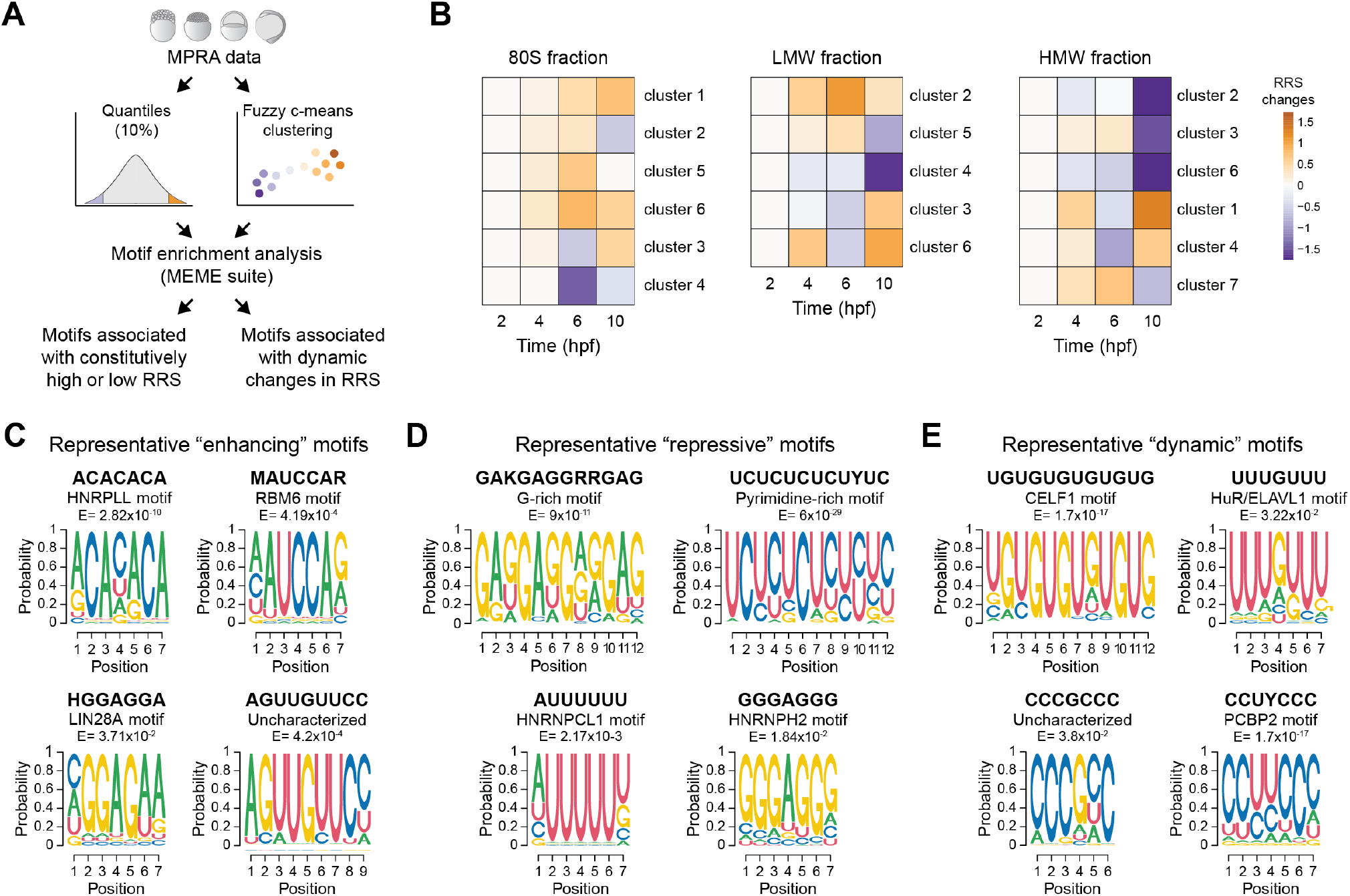
The *in vivo* MPRA identifies 5’ UTR *cis*-regulatory motifs. (A) Schematics of the strategy used for grouping 5’ UTRs for motif-based sequence enrichment analysis based on RRS values during embryogenesis. (B) Clusters derived from fuzzy-c means clustering of mean log2 RRS values for the 80S, low molecular weight (LMW) and high molecular weight (HMW) fractions. The colour of each square represents the difference in cluster means (log2 RRS values) for each developmental time-point normalized to 2 hpf. Only clusters containing 5’ UTRs with membership score ≥ 0.7 are represented. Note, that these clusters differ from the ones in Figure 2. (C) Representative motifs enriched in 5’ UTRs that are consistently found in the upper quantile (top 10%, “enhancing”) or (D) the lower quantile (bottom 10%, “repressive”) throughout the developmental time-course, or (E) enriched in 5’ UTRs with high cluster membership scores (≥ 0.7) (“dynamic”). See STAR Methods, Figure S4 and Table S4.

Next, we focused on identifying motifs enriched in 5′ UTRs that display dynamic changes in RRS throughout early embryogenesis (Figure 4A). For each fraction, we performed unsupervised soft clustering analysis (Futschik & Carlisle, 2005; Kumar & M, 2007) (see STAR Methods for cluster analysis parameters) of RRS values across developmental time (2, 4, 6 and 10 hpf) for each fraction (80S, LMW and HMW), which identified clusters of 5′ UTRs that display coordinated changes in RRS across early embryogenesis (Figure 4B; Table S3). We manually grouped clusters that showed qualitatively similar dynamics (Figures S4B and S4C) and ran motif enrichment analysis using the MEME tool suite (Bailey *et al*., 2015). This analysis returned an additional 25 motifs associated with temporal RRS dynamics, of which 14 are *de novo* predicted motifs (Table S4).

We categorized motifs as “enhancing”, “repressive” or “dynamic” based on whether they were enriched in the top 10% RRS quantiles, bottom 10% quantiles, or in any of the dynamic-based clusters (Figures 4C-E and S4D-F; Table S4). Importantly, and having in mind that each motif enrichment analysis was performed independently, motifs identified in opposing quantiles are largely non-overlapping. Moreover, similar motifs were commonly enriched across different fractions. For example, CA-rich motifs such as ACACACA or similar derivatives (MAUCCAR or AMAWACA) were consistently enriched in the top quantile of all fractions (Figure 4C and S4D; Table S4). In contrast, G- and GC-rich motifs (such as GAKGAGGRRGAG and GMGCGCKCGSYC) and pyrimidine-rich motifs (such as UCUCUCUCUYUC and CCCUCUCYCYCY) were enriched in 5′ UTRs associated with poor ribosome recruitment (Figure 4D and S4E; Table S4). For some cases, the motif identified was enriched in only one of the fractions, as was the case for “enhancing” motifs HGGAGGA (top 10% 80S fraction), ACUUCCGG (top 10% LMW fraction) and AGUUGUUCC (top 10% HMW) (Figures 4C and S4D), or “repressive” motifs AUUUUU (bottom 10% 80S fraction), GGGAGGG (bottom 10% LMW) and CAGAAGAGCAGC (bottom 10% HMW) (Figures 4D and S4E).

Among the “dynamic” motifs, UGU-rich motifs enriched in 5′ UTRs that display a decrease in RRS at the bud (10 hpf) stage stood out, namely UGUGUGUGUGUG, UGUKURUKU and UUUGUUU, as each was independently identified in distinct clusters that display analogous RRS dynamics (Figure 4E; Table S4). Shorter motifs (5 to 7 nt-long) may also contribute to the temporal regulation of translation initiation frequency and consequently transcript ribosome load. For example, the motif CCCGCC (80S cluster 1) is associated with monosome recruitment that gradually increases during gastrulation (from 6 hpf onwards) whereas the motif CCUYCCC (LMW cluster 3) is enriched in 5′ UTRs with similar temporal behavior but that lead to the recruitment of 2 to 4 ribosomes per transcript. In contrast, motifs with GA dinucleotides (e.g., GAGAGARAGAGA, AGAGAAA and GAAGAAG) are associated with dynamic recruitment to polysomal fractions throughout the developmental time-course (Figure S4F; Table S4). Note that some motifs with high similarity can be categorized as both enhancing/repressive and dynamic (e.g., the “enhancing” ACACACA motif and the “dynamic” CACACACACACA motif) (Table S4). In summary, we identified 86 motifs (5-12 nt-long) associated with distinct translation dynamics, 44 of which are newly discovered motifs with putative roles in translation initiation control.

### DaniO5P predicts the effect of 5’ UTR sequences on translation dynamics

Our analyses identified individual 5′ UTR features that covary with RRS, but they do not integrate complex interactions such as the relationship between 5′ UTR length, uORF number and motif representation. To address this, we used the *in vivo* MPRA data to train predictive models based on convolutional neural networks (CNNs). Following our previous modeling work in human cells (Sample *et al*., 2019), we chose to summarize the translation state of every reporter at each time-point via the mean ribosome load (MRL), calculated by taking the average of transcript abundances in the 80S, LMW, and HMW fractions weighted by their approximate mean ribosome number (Figure 5A) (see STAR Methods for calculation). As with RRS, we found MRL to be highly dependent on 5’ UTR length. In fact, a simple "length model" based on a quadratic polynomial at each time-point explained 64-85% of the MRL variance (Figure 5C and S5A; Table S4). To capture how sequence features beyond length may influence MRL, we trained an ensemble of 10 CNNs to predict MRL residuals (observed - length model prediction) at all four time-points simultaneously given an arbitrary 5’ UTR sequence (Figure 5B) (STAR Methods). Remarkably, CNN predictions explained 41-52% of the remaining variation in sequences held out from training, and the combined length & CNN model accounted for 79-93% of the overall MRL variation (Figure 5C). We termed the combined length & CNN model *Danio* Optimus 5-Prime (DaniO5P). DaniO5P captured observed MRL dynamics, explaining, for example, 73% of the variation in MRL differences at 10 versus 2 hpf, whereas the length model alone could only explain 51% of the variation (Figure 5C and S5B). For example, the predicted MRL for reporters bearing the 5′ UTR of transcripts usp12a, crygm2d13, unc119b, and bzw1b more closely matched the measured translation patterns during embryogenesis when integrating both length and sequence features in the model predictions (Figure 5D). These results indicate that 5′ UTR length and sequence element grammar cooperate to fine tune translation initiation.

**Figure 5.**
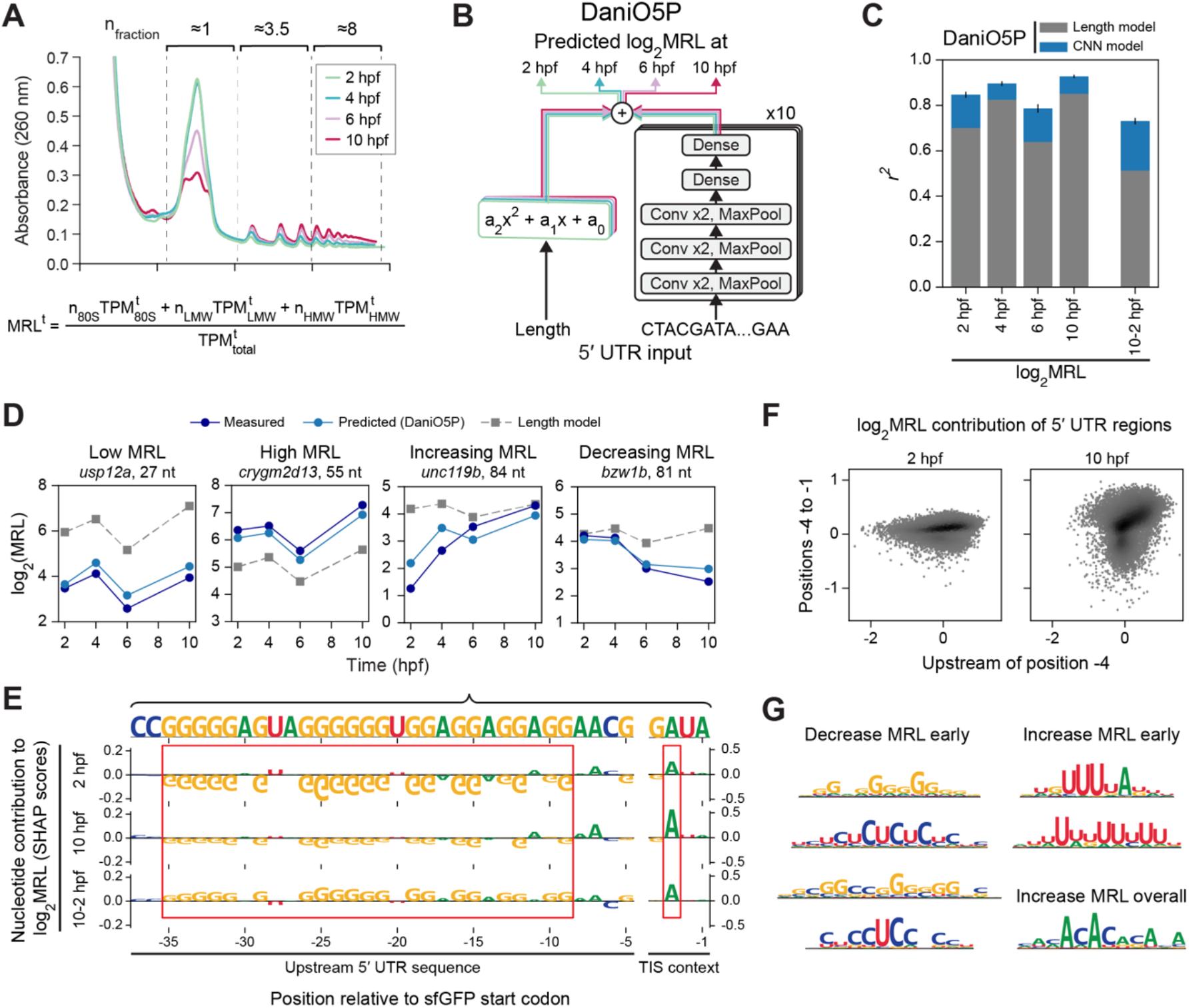
DaniO5P reveals sequence determinants of translation dynamics. (A) MRL calculation based on polysome profiling. (B) Predictive DaniO5P model schematic. log2(MRL) predictions at 2, 4, 6, and 10 hpf result from the combination of a second-order polynomial on the 5’ UTR length and an ensemble of 10 convolutional neural networks that incorporate sequence features. (C) Prediction performance separated by contributions of the length and sequence model, for all four time-points and for the difference between 10 and 2 hpf. (D) Examples of translation dynamics in 5’ UTRs that substantially differ from what would be expected from their length alone, showing that CNN model predictions can account for these differences. (E) Nucleotide contributions (SHAP values, see STAR Methods) of the 5’ UTR of transcript unc119b at 2hpf, 10hpf, and their difference. G repeats are shown to decrease MRL at earlier time-points whereas an A at position −3 increases MRL at later time-points, both contributing to the overall MRL increasing dynamics. Note that the y axis scale was adjusted differently for the TIS context region and the rest of the 5’ UTR sequence. (F) Sum of nucleotide contributions within the TIS context region (positions −4 to −1) and outside (upstream of −4) for every sequence in the MPRA. Sequences outside the TIS context region contribute more strongly at earlier time-points, whereas the TIS context region has a larger contribution later. (G) Examples of motifs extracted from the DaniO5P model, along with a description of their average contribution to MRL. See also Figure S5 and Table S4.

To recover sequence features learned by DaniO5P, we calculated, for every sequence in the MPRA, the contribution of each nucleotide to MRL predictions at each developmental time-point (Table S4; STAR Methods). Contribution scores recapitulated expected sequence features such as upstream AUGs, which contribute negatively to MRL at all time-points (Figure S5C and G). The predicted impact of the region immediately upstream of the sfGFP start codon (positions −4 to −1), whose effect depends on their nucleotide composition (Figure S5C-F), is consistent with the observed TIS sequence context (Figure 3D). Interestingly, we found the effect of some features to change throughout embryogenesis. As an example, long stretches of Gs in the 5′ UTR of unc119b negatively contribute to MRL at 2 hpf, but this effect is diminished by the end of gastrulation (10 hpf) (Figure 5E). In contrast, the presence of an adenosine (A) at position −3 of the main sfGFP ORF has a stronger enhancing contribution at 10 hpf than at 2 hpf. More generally, across all sequences, the immediate TIS context of the sfGFP ORF (positions −4 to −1) showed a stronger contribution to MRL at later stages of embryogenesis (10 hpf compared to 2 hpf), whereas nucleotides outside of this region (position −5 and upstream) contribute more strongly at earlier stages (Figure 5F).

Finally, we identified specific sequence motifs learned by DaniO5P. We extracted position weight matrices (PWMs) that maximally activate convolutional filters, clustered them to avoid redundancy, and filtered based on their reproducibility across independently trained models. We then estimated their contribution to MRL by averaging the nucleotide contributions within motif occurrences across all MPRA sequences (STAR Methods). This analysis resulted in 16 reproducible motifs which are predicted to contribute to changes in MRL throughout embryogenesis (pyrimidine-rich motifs, G- and GC-rich motifs, U-rich motifs), or to overall enhancement (CA-rich motifs) or dampening of MRL (uAUGs) (Figures 5G and S5G; Table S4). The most inhibitory uAUGs have a strong TIS context, that is, with A/G at position −3 and/or G at position +4 (Figure S5G). Of note, DaniO5P predicts U-rich motifs as most enhancing during early embryogenesis (2 and 4 hpf) whereas CU-rich motifs become less inhibitory at later stages (10 hpf) (Figure S5G; Table S4), agreeing with findings reported by a complementary 5′ UTR MPRA method (Strayer *et al*., 2023).

Pair-wise comparison of all motifs identified by DaniO5P or by sequence enrichment analysis using the Tomtom motif comparison tool pinpoints motifs with significant sequence similarity (p-value < 0.05) (Figure S6A; Table S4). Motifs with high similarity score (p < 1×10^−4^) generally display analogous effects on predicted MRL and measured RRS (Figures 4C-E, S4D-E and S5G). For example, model and RRS analyses indicate that pyrimidine-rich motifs and G-rich motifs are inhibitory and that UGU-rich motifs are associated with a decrease in ribosome recruitment during embryogenesis. Similarly, both analyses pinpoint CA-rich motifs as enhancing that can be associated with an increase in ribosome recruitment as embryogenesis progresses. Thus, DaniO5P is remarkably predictive of 5′ UTR activity and provides a powerful approach to define the interaction of distinct 5′ UTR features on translation initiation regulation.

### Switching 5’ UTR isoforms display different translation initiation capabilities during the maternal-to-zygotic transition

During the zebrafish MZT, differential TSS utilization can result in maternally deposited and zygotically expressed transcript isoforms that bear different 5′ UTR sequences but still encode the same gene product (Figure 6A). We hypothesized that switching 5′ UTRs could provide an additional layer to gene expression regulation by modulating transcript translation. Analysis of differential TSS utilization identified cases of maternal-zygotic isoform 5′ UTR switching events (Figure 6A; Tables S1 and S5), as has been described previously (Haberle *et al*., 2014; Nepal *et al*., 2013). Transcript 5′ UTR isoforms display a similar length distribution (median = 126 nt-long; Figure S6B), indicating that switching does not favor shortening nor lengthening of the 5′ UTR of zygotically expressed isoforms. TSS switching is bi-directional (Haberle *et al*., 2014) and tends to alter 5′ UTR length by a narrow size window (Figure S6C; Table S5), with a mean length shift of 37 nts between maternal and zygotic transcript isoform pairs.

**Figure 6.**
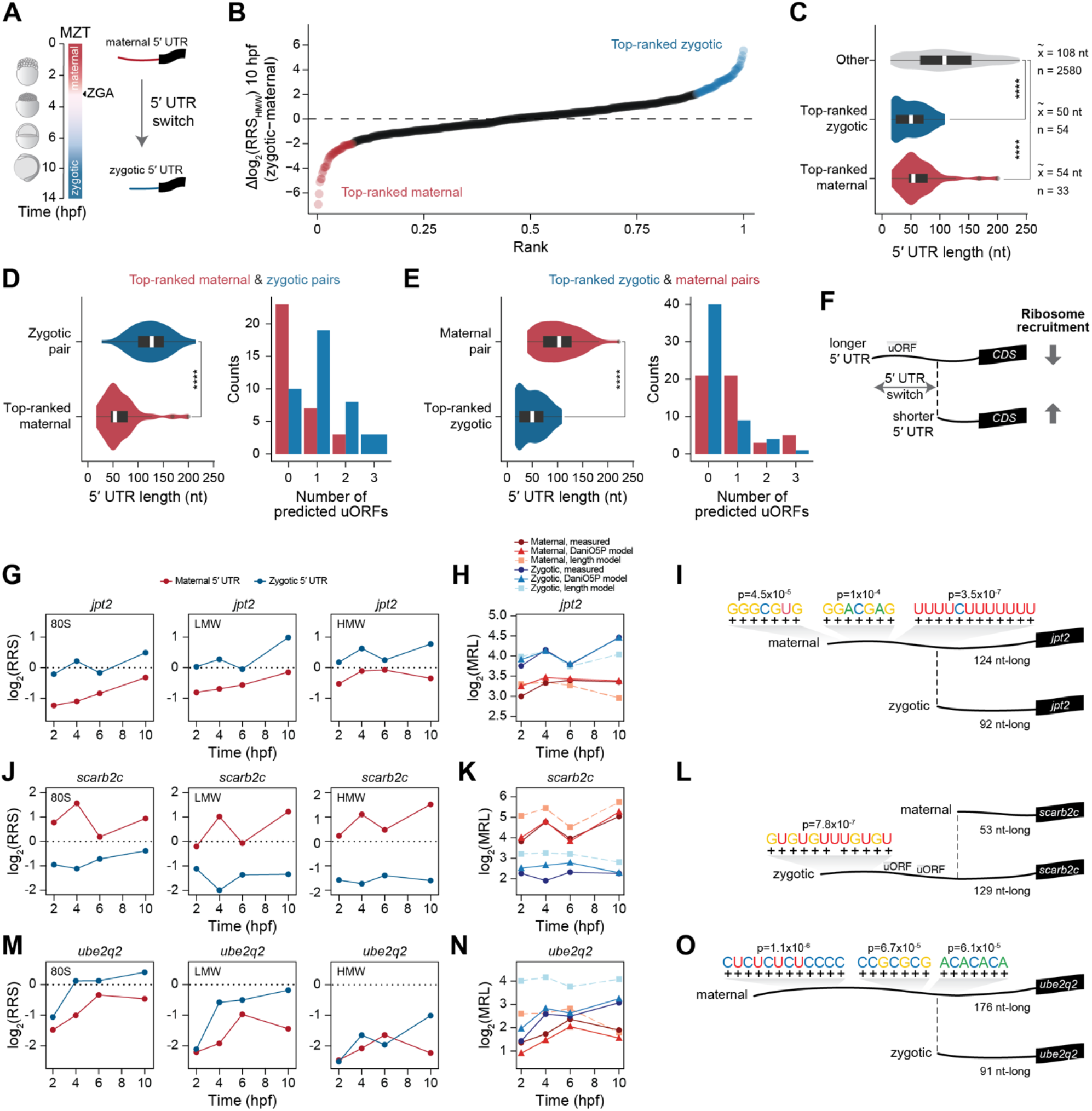
Switching 5’ UTR isoforms display different translation initiation capabilities during the maternal-to-zygotic transition. **(A)** 5’ UTR isoform switches arising during the maternal-to-zygotic transition (MZT). Maternal isoforms are deposited in the mother’s egg and later, during zygotic genome activation (ZGA), a longer or shorter zygotic 5’ UTR isoform is expressed. (B) Difference in mean log2 transformed RRS values for the HMW fraction at 10 hpf between zygotic and maternal switching 5’ UTR isoforms pairs. Cases for which the maternal 5’ UTR isoform leads to higher ribosome recruitment ((Δlog2(RRSHMW)) ≤ −2) are marked as “top-ranked maternal” (n = 33) and for which the zygotic leads to higher ribosome recruitment (Δlog2(RRSHMW) ≥ 2) are marked as “top-ranked zygotic” (n = 54). Only 5’ UTRs shorter than 239 nts were considered. (C) Violin plots of the length distribution of top-ranked maternal (n = 33), top-ranked zygotic (n = 54) and other (n = 2,580) switching isoforms. Center line of boxplot represents median, box limits represent the upper and lower quartiles. **** p-value < 0.001, two-sample rank sum Wilcoxon test. (D) Violin plot of the length distribution and bar blot of the number of uORFs for top-ranked maternal isoforms and respective zygotic pairs, and (E) for top-ranked zygotic isoforms and respective maternal pairs. Center line of boxplot represents median, box limits represent the upper and lower quartiles. **** p-value < 0.001, two-sample rank sum Wilcoxon test. (F) Isoform switching events that shorten 5’ UTR length and remove uORFs lead to increased ribosome recruitment. (G, J, M) Line plots of mean log2 transformed RRS values for 80S, LMW and HMW fractions for examples of maternal and zygotic switching 5’ UTR pairs (transcripts jpt2, scarb2c and ube2q2) with distinct translational dynamics and (H, K, N) corresponding line plots of measured and predicted log2 transformed MRL values for length and DaniO5P (length + CNN) models. (I, L, O) Schematics depicting motif sequences matches and respective p-values. See also Figure S6, S7, Table S4 and S5.

To determine whether maternal and zygotic 5′ UTR isoforms confer distinct translation initiation capabilities, we restricted our analysis to switching 5′ UTRs that were assayed in the MPRA as full uninterrupted sequences (n = 1,405 out of 2,721, that range between 15-238 nts). We calculated the difference in RRS (Δlog_2_(RRS_HMW_)) for the HMW fraction at the end of gastrulation (bud stage, 10 hpf) between zygotic and maternal switching isoforms. By ranking Δlog_2_(RRS_HMW_) values, we find that 5′ UTR isoform switching leads to up to 2 orders of magnitude differences in ribosome recruitment to the mRNA reporter (n = 507 5′ UTR pairs; Figure 6B and Table S5). Switching resulted in higher translatability of the reporter bearing the maternal 5′ UTR (Δlog_2_(RRS_HMW_) < 0) or the zygotic 5′ UTR (Δlog_2_(RRS_HMW_) > 0), indicating that isoform switching does not impose a unidirectional effect on translation initiation capability.

The length of the 5′ UTR and uORFs impact ribosome recruitment (Figures 3 and 5). We therefore selected 5′ UTR pairs that display largest differences in ribosome recruitment capability (absolute Δlog_2_(RRS_HMW_) ≥ 2) between zygotic and maternal isoforms and determined their 5′ UTR length and number of predicted uORFs. For 5′ UTR pairs with largest differences in RRS, there were 33 5′ UTRs for which the maternal isoform was better initiated (Δlog_2_(RRS_HMW_) ≤ −2, “top-ranked maternal”) and 54 5′ UTRs for which the zygotic isoform was better initiated (Δlog_2_(RRS_HMW_) ≥ 2, “top-ranked zygotic”) (Figure 6B; Table S5). As anticipated, top-ranked isoforms were shorter than all other 5′ UTR variants regardless of it being of maternal or zygotic origin (p-value < 0.001, Wilcoxon rank sum test; Figure 6C). Moreover, when considering switching pairs, we observe that the 5′ UTR length of the top-ranked maternal isoform was significantly shorter than its zygotically-expressed counterpart and the same was true for top-ranked zygotic isoforms and their maternal isoform pairs (Figures 6D and E). Similarly, top-ranked 5′ UTR isoforms had fewer uORFs than their zygotic or maternal variant pairs (Figure 6D and E), implying that TSS switching can result in the exclusion of inhibitory uORFs. We made similar observations for isoforms arising from switching events throughout ZGA (ZGA and postZGA 5′ UTR switching pairs, n = 330) (Figure S6B-G; Table S5). These data demonstrate that modulation of 5′ UTR length and uORF number by alternative TSS switching is a simple yet powerful mechanism to regulate translation during embryogenesis (Figure 6F).

We wondered if 5′ UTR switching also alters motif grammar. First, we used MAST (Bailey & Gribskov, 1998) to search for the occurrence of motifs identified by the MPRA (Figure 4 and Table S4) in switching UTRs. MAST scored motif matches (p < 10^-4^) in 228 of the 2,721 switching 5′ UTRs (Table S4). By manually parsing sequences, we identified several cases where switching alters motif occurrence in the 5′ UTR. For example, maternal and zygotic jpt2 5′ UTR isoforms display distinct RRS patterns during early embryogenesis, and the zygotic 5′ UTR isoform leads to preferential ribosome recruitment at the bud stage (10 hpf) (Figure 6G). To determine how much of the diverging translational behavior between 5′ UTR isoforms can be explain by sequence features, we turned to the DaniO5P model. DaniO5P more accurately predicts the MRL of maternal and zygotic 5′ UTR isoforms than the length model alone (Figures 6H, K, N and S7A-D), supporting that changes to 5′ UTR motif grammar contribute to their differential translatability. For example, switching between maternal and zygotic jpt2 5′ UTRs shortens its length by 32 nts and removes a stretch of sequence matching three distinct motifs (GGGCGUG, p-value = 4.5×10^−5^; GGACGAG, p-value = 1×10^−4^; UUUUCUUUUUUU, p-value = 3.5×10^−7^) (Figure 6I). Accordingly, sequence contribution scores point at the importance of these nucleotide stretches to the final output of DaniO5P predictions (Figure S6H).

Similarly, the maternal 5′ UTR isoform of scarb2c leads to reporter ribosome recruitment throughout the time-course, whereas the zygotic 5′ UTR isoform does not, and model predictions that integrate motif grammar better predict their regulatory capabilities (Figures 6J and K). Isoform switching of scarb2c leads to the inclusion of a 12 nt stretch matching a UGU-rich motif (p-value = 7.8×10^−7^) and two uORFs in the zygotic 5′ UTR (Figure 6L and S6I). Both the motif enrichment analysis (Table S4) and the DaniO5P model (Figure S5G) pinpointed GU-rich motifs as generally inhibitory. For ube2q2, switching results in removal of a pyrimidine-rich motif (CUCUCUCUCCCC, p-value = 1.1×10^−6^) and a GC-rich motif (CCGCGCG, p-value = 6.7×10^−5^), that are predicted as important contributing nucleotides stretches (Figure S6J), as well as the disruption of a CA-rich motif (ACACACA, p-value = 6.1×10^−5^) (Figure 6O). Interestingly, we also find cases where one of the 5′ UTR isoforms promotes preferential 80S recruitment (Figure 6M, S7A and S7B). These and additional examples (Figure S7) showcase how length alone cannot fully explain the regulatory capacity of 5′ UTR sequences: the maternal jpt2 5′ UTR and the zygotic scarb2c 5′ UTR are of comparable length (124 and 129 nts, respectively), and so are the zygotic 5′ UTR isoforms of jpt2 and ube2q2 (92 and 91 nts, respectively), yet their translational behaviors are distinct. Our results suggest that TSS switching contributes to regulating the translation of transcript isoforms by modulating 5′ UTR length and motif grammar.

## Discussion

Our comprehensive *in vivo* characterization of 5′ UTRs and their role in translation initiation provides six main conclusions. First, 5′ UTR sequences modulate the temporal dynamics of translation during embryogenesis (Figure 2). Second, the MPRA reveals the regulatory effects of RNA structures, uORFs and sequence context on ribosome recruitment. Third, 5′ UTR length is a major determinant of translation initiation *in vivo* (Figure 3). Fourth, known conserved and novel uncharacterized 5′ UTR *cis*-acting motifs are associated with differential ribosome recruitment during embryogenesis (Figures 4 and 5). Fifth, shifts in 5′ UTR length and motif grammar alter translation initiation during the zebrafish MZT (Figure 6). Finally, the deep learning model DaniO5P predicts ribosome recruitment solely based on 5′ UTR sequence (Figures 5 and 6) and provides a powerful tool to dissect the regulatory logic of 5′ UTR elements.

### Regulatory features of 5′ UTRs acting during embryogenesis

Our work provides the largest-scale characterization to date of the regulatory capacity of endogenous 5′ UTR sequences during zebrafish embryogenesis. Translational control confers a means to rapidly regulate protein synthesis during the fast-paced process of embryogenesis (Harnett *et al*, 2022; Ozadam *et al*, 2023; Teixeira & Lehmann, 2019; Xiong *et al*, 2022). While it has long been appreciated that 5′ UTR sequences are the first point of control for regulating protein synthesis, the contribution of 5′ UTRs to determining when and how efficiently an mRNA is translated during developmental transitions has been unclear.

Our 5′ UTR MPRA recapitulates the widespread negative impact of RNA structure and uORFs on ribosome recruitment *in vivo* (Calvo *et al*., 2009; Chew *et al*., 2016; Cuperus *et al*., 2017; Johnstone *et al*., 2016; Leppek *et al*., 2018; Lin *et al*, 2019; May *et al*., 2023; Niederer *et al*., 2022; Sample *et al*., 2019; Zhang *et al*, 2019) (Figure 3C), and shows that canonical uAUGs with a strong TIS sequence context (purine at position −3 and/or G at +4) are most detrimental for ribosome recruitment (Figure S5G). The data reaffirms the functional importance of the TIS consensus context AAAC<u>AUG</u> for efficient TIS recognition (Giess *et al*., 2020; Grzegorski *et al*., 2014; Kozak, 1986b) in zebrafish and revealed that the presence of a U nucleotide at positions −4 to −2 upstream of the TIS is particularly detrimental for translation initiation (Figure 3D). Our results suggest that their negative impact on translation initiation is conserved across vertebrates, since mutagenesis of the TIS to 4Us (positions −5 to −1) abolished protein synthesis from a reporter plasmid in mammalian cells (Kozak, 1986b), and Us at positions −3 and −2 are universally absent in the consensus TIS of annotated CDSs (Hernandez *et al*., 2019; Shabalina *et al*, 2004). A recent method for immunopurification of epitope tagged nascent peptides driven by translation initiation of a reporter 5′ UTR pool with random nucleotides upstream of the TIS also pinpointed A/C nucleotides as enhancing and U at position −3 as particularly detrimental for initiation in zebrafish embryos (Strayer *et al*., 2023). Interestingly, the quantitative DaniO5P model indicates that positions −4 to −1 of the TIS context are less deterministic of ribosome recruitment at earlier stages of embryogenesis (2 hpf) (Figure 5F). It is conceivable that the limited availability of free ribosomes at early stages of zebrafish embryogenesis (Leesch *et al*., 2023) amplifies the regulatory effects of other 5′ UTR sequence features on translation initiation, and perhaps reflects prominent RBP-mediated regulation at early stages of development (Despic *et al*, 2017).

Notably, we find that the 5′ UTR length has a major impact on translation initiation *in vivo* (Figures 3B and 5C). Once the PIC is engaged with the mRNA 5′ end, scanning starts immediately and continues in a processive manner throughout the 5′ UTR (Giess *et al*., 2020; Wang *et al*., 2022). In a scenario where the PIC remains tethered to the 5′ cap while scanning the 5′ UTR, which would block the entry of a new PIC during ongoing scanning, the length of the 5′ UTR would be limiting for translation efficiency (Bohlen *et al*., 2020; Chappell *et al*., 2006; Shirokikh & Preiss, 2018). Thus, our observations are consistent with scanning being mainly cap-tethered in zebrafish embryos.

### A path toward the discovery of 5′ UTR-binding RBPs

Our study defines a set of short sequence motifs enriched in 5′ UTRs that confer distinct amplitudes to ribosome recruitment throughout early embryonic development (Figures 4 and 5). Eukaryotic mRNAs interact with RBPs throughout their life-cycle. RBPs generally make contacts with 3-6 consecutive RNA bases and some RBPs favor longer spaced “bipartite” motifs (Afroz *et al*, 2015; Auweter *et al*, 2006; Dominguez *et al*., 2018). RBPs and their sequence specificities are highly conserved (Beckmann *et al*, 2015; Despic *et al*., 2017; Gebauer *et al*, 2021; Gerstberger *et al*., 2014; Matia-Gonzalez *et al*, 2015; Ray *et al*., 2013; Sysoev *et al*, 2016; Wessels *et al*, 2016), but their roles in embryogenesis are poorly understood. Many of the motifs identified by our MPRA match the consensus sequence of known RBPs in other species (Ray *et al*., 2013), and homologs of those RBPs are expressed during zebrafish embryogenesis (Figure S4G).

Among the motifs enriched in 5′ UTRs, we find for example the consensus binding sequence for the translational regulators IGF2BP2 (AMAWACA motif) and PCBP2 (CCUYCCC motif). IGF2BP2 is a mouse embryonic RBP shown to directly bind the 5′ UTR of target transcripts (Dai *et al*, 2011; Dai *et al*, 2015; Nielsen *et al*, 1999), and PCBP2 is a translational inhibitor acting via 5′ UTR-binding involved in mouse erythropoietic differentiation (Ji *et al*, 2021; Smirnova *et al*, 2019). Other motif-matching RBPs have been described to bind the 3′ UTR of target transcripts to regulate translation, such as Lin28A (HGGAGAA motif) and HuR (UUUUUUU and UUUGUUU motifs). Lin28A is expressed in human stem cells (Yu *et al*, 2007) and regulates the translation of a subset of mRNA targets (Cho *et al*, 2012; Polesskaya *et al*, 2007; Qiu *et al*, 2010; Xu & Huang, 2009; Xu *et al*, 2009). While most mRNA Lin28A-binding sites have been identified in exonic and 3′ UTR regions (Wilbert *et al*, 2012), its interaction with translation initiation complexes including the 5′ cap-binding protein (Polesskaya *et al*., 2007) and 5′ to 3′ RNA helicases (Jin *et al*, 2011; Parisi *et al*, 2021) raises the possibility that the regulation of translation initiation via the 5′ UTR of a few select mRNA targets may involve the Lin28A RBP. HuR required for normal mouse embryogenesis (Han *et al*, 2022; Katsanou *et al*, 2009) and regulates the temporal association of functionally related mRNAs in actively translating polysomes during mouse neurogenesis (Kraushar *et al*, 2014; Popovitchenko *et al*, 2016). HuR mostly interacts with mRNAs via the 3′ UTR, but motifs can be also be located within the 5′ UTR of target transcripts (Lopez de Silanes *et al*, 2004; Mukherjee *et al*, 2011). We postulate that the motifs identified by the MPRA might provide a platform for RBP-mediated regulation of translation initiation during embryogenesis.

Notably, the majority of enriched RBP motifs match those of proteins with roles in pre-mRNA splicing, such as SR proteins and heterogeneous nuclear ribonucleoproteins (hnRNPs). In *Drosophila*, the splicing factor Sex-lethal (SXL) binds U-rich motifs in the 5′ UTR of the msl-2 mRNA to block scanning (Bashaw & Baker, 1997; Beckmann *et al*, 2005; Gebauer *et al*, 1998; Kelley *et al*, 1997) and ensure dosage compensation during development (Conrad & Akhtar, 2012). These and other studies (Long & Caceres, 2009; Maslon *et al*, 2014; Palangat *et al*, 2019; Sanford *et al*, 2004; Ueno *et al*, 2019) show that splicing factors can play splicing-independent regulatory roles in translation regulation. Our findings are also in agreement with previous RNA interactome capture experiments that showed that RBPs that dynamically bind to cytoplasmic mRNAs during the fly and zebrafish MZT are enriched for proteins involved in mRNA splicing (Despic *et al*., 2017; Sysoev *et al*., 2016). These observations raise the possibility that splicing factors may be playing broad uncharacterized roles in translational control via 5′ UTR-binding during early zebrafish embryogenesis.

### Regulatory potential of 5′ UTR isoforms

Our study demonstrates that the regulatory potential of 5′ UTR sequences during embryogenesis can be expanded by the expression of 5′ UTR isoforms. We included 5′ UTR variants in our MPRA arising from TSS switches throughout zebrafish early embryogenesis and found that 5′ UTR switching alters their capability to recruit ribosomes (Figure 6). Our *in vivo* findings extend previous studies in mammalian cells and yeast that reported that longer 5′ UTR isoforms are associated with lower translation (Blair *et al*., 2017; Floor & Doudna, 2016; Rojas-Duran & Gilbert, 2012; Wang *et al*., 2016; Wong *et al*., 2016) and that the inclusion of inhibitory uORFs in extended 5′ UTRs reduces translation of the main open reading frame (Capell *et al*, 2014; Chen *et al*, 2017; Cheng *et al*., 2018; Hollerer *et al*, 2019; Tresenrider *et al*., 2021). MPRA measurements and DaniO5P predictions indicate that simply shortening the 5′ UTR sequence provides an elegant and efficient mechanism to modulate protein synthesis capability. Finally, zebrafish switching 5′ UTR variants can display different motif grammar, and DaniO5P indicates that changes to motif composition impacts their translational effect. Interestingly, cancer cells take advantage of TSS switches to expose or mask motifs to promote translation of a cohort of growth-promoting transcripts (Weber *et al*, 2023). Moreover, two studies have reported 5′ UTR isoform-specific translation by binding of an RBP to one of the 5′ UTR isoforms but not the other (Aeschimann *et al*, 2017; Popovitchenko *et al*, 2020). Thus, differential RBP binding to 5′ UTR variants might contribute to isoform-specific translational regulation to coordinate protein synthesis during zebrafish embryogenesis.

Altogether, these results lay the foundation for future work that will determine the identity and roles of RBPs that interact with the motifs described in this study. Our work highlights the regulatory diversity within vertebrate 5′ UTRs, and shows the power of combining sequence-based analyses with deep learning to disentangle the impact of 5′ UTR features on translation initiation *in vivo.* We anticipate that DaniO5P’s predictive power will enable the design of synthetic 5′ UTRs (Castillo-Hair *et al*., 2023; Castillo-Hair & Seelig, 2022) with desired translational behaviors during zebrafish embryogenesis.

### Limitations of study

The cloning and sequencing strategy we employ here requires the presence of a common adaptor at the most 5′ end of the reporter. Thus, the regulatory impact of motifs with 5′ end positional dependent activity, such as the 5′ terminal oligopyrimidine (TOP) motif (Meyuhas & Kahan, 2015), cannot be recovered by the MPRA. We observed that some of the switching events result in the establishment of a 5′ TOP motif at the most 5′ end of the shorter 5′ UTR isoform (Figure S7E). The functional consequence of such switching events during embryogenesis warrants further investigation. Case studies of endogenous transcripts have shown that RBP binding sites in 3′ UTRs can mask the regulatory effects from sites in the 5′ UTRs (and vice-versa) (Theil *et al*, 2018). Thus, it is possible that the regulatory effects of individual 5′ UTRs recovered by the MPRA may differ from its regulatory potential in the context of the endogenous transcript. Our work does not consider the presence of epitranscriptomic marks (Seo & Kleiner, 2021) in the 5′ UTR, such as *N*^6^-methyladenosine (m^6^A), which was shown to promote cap-independent translation initiation (Meyer *et al*, 2015).

## Acknowledgments

We are very grateful to Dr. Sunil Shetty for polysome profiling training and for assistance collecting polysome fractions. We thank Dr. Andrea Pauli and her lab for advice regarding polysome profiling buffer composition for zebrafish embryos. We thank Dr. Matthias Muhar, Dr. Júlia Batki and Dr. Annika Nichols for advice throughout the project and for critical reading of the manuscript. We thank the Schier lab and the Basel RNA Club for input and discussions, and the Biozentrum Core Facilities for technical support. We thank Alba Aparicio Fernandez and Rita Gonzalez Dominguez for support with fish husbandry. High-throughput sequencing was performed at the Genomics Facility Basel. This project has received funding from the European Union’s Horizon 2020 research and innovation programme under the Marie Skłodowska-Curie grant agreement No 898218 and from the EMBO long-term fellowship ALTF 691-2019 awarded to M.M.R-P., from the Allen Discovery Center for Cell Lineage Tracing and an ERC Advanced grant to A.F.S., and from NIH Award R33CA255893 and NSF Award and NSF Award 2021552 to G. S.

## Author contributions

Study conceptualization M.M.R-P. and A. F. S. Design, methodology, experimental investigation and formal analysis, M.M.R-P. Generation of the CNN model and related analyses, S.M.C-H. Visualization – figure generation, M.M.R-P. and S.M.C-H. Writing – original draft, M.M.R-P. and S.M.C-H. Writing – reviewing & editing, M.M.R-P., S.M.C-H, A.F.S. and G.S. Supervision and Funding Acquisition, M.M.R-P., A.F.S. and G.S.

## Supplemental Figures

**Figure S1.**
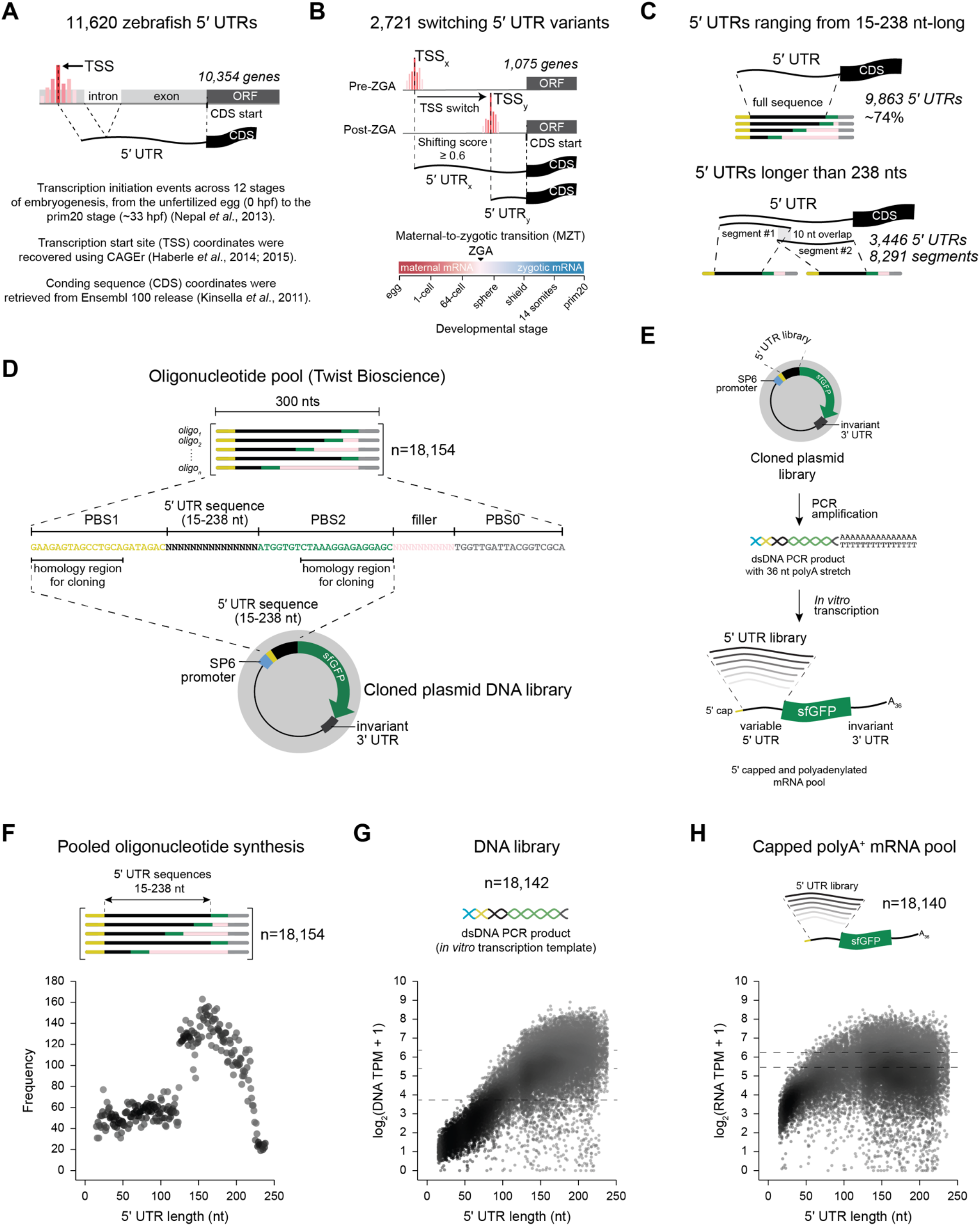
– Schematics of the approach for recovering 5′ UTR sequences, related to Figure 1. 5′ UTR sequences of 10,354 genes expressed throughout embryogenesis were annotated by recovering the exonic sequence between the dominant transcription start site (TSS) and the annotated coding sequence (CDS) start site; introns were excluded. (B) 5′ UTR isoform sequences arising from differential TSS utilization (shifting score ≥ 0.6, p-value ≤ 0.01, K-S test FDR ≤ 0.01) throughout the maternal-to-zygotic transition were considered. (C) Of the 5′ UTRs tested, 9,863 correspond to the full uninterrupted endogenous sequence (∼74% of all 5′ UTRs); the 3,446 remaining ones longer than 238 nt were split into segments with overlapping 10 nts, amounting to a total of 8,291 split 5′ UTRs. (D) The oligonucleotide pool was commercially synthetized by Twist Bioscience, and amplified using a two-step PCR approach via primer binding sites PBS0, PBS1 and PBS2 followed homologous recombination-mediated cloning. (E) Strategy used for generation of a dsDNA template compatible with pooled *in vitro* transcription of a capped and polyadenylated 5′ UTR mRNA library. (F) Scatter plot representing the length (x axis) and the number of 5′ UTR sequences of each length (y axis) of the *in silico* designed 5′ UTR library. 5′ UTR sequences assayed range between 15 and 238 nt in length (black color), and include a 22 nt-long common upstream adaptor (yellow color) that is not considered in the total length of the 5′ UTR. (G) Scatter plot representing the length (x axis) and the abundance of 5′ UTR sequences of each length (y axis) of the DNA library PCR product used as template for *in vitro* transcription. (H) Scatter plot representing the length (x axis) and the abundance of 5′ UTR sequences of each length (y axis) of the *in vitro* transcribed mRNA library used for embryo injections. See also STAR Methods and Table S1.

**Figure S2.**
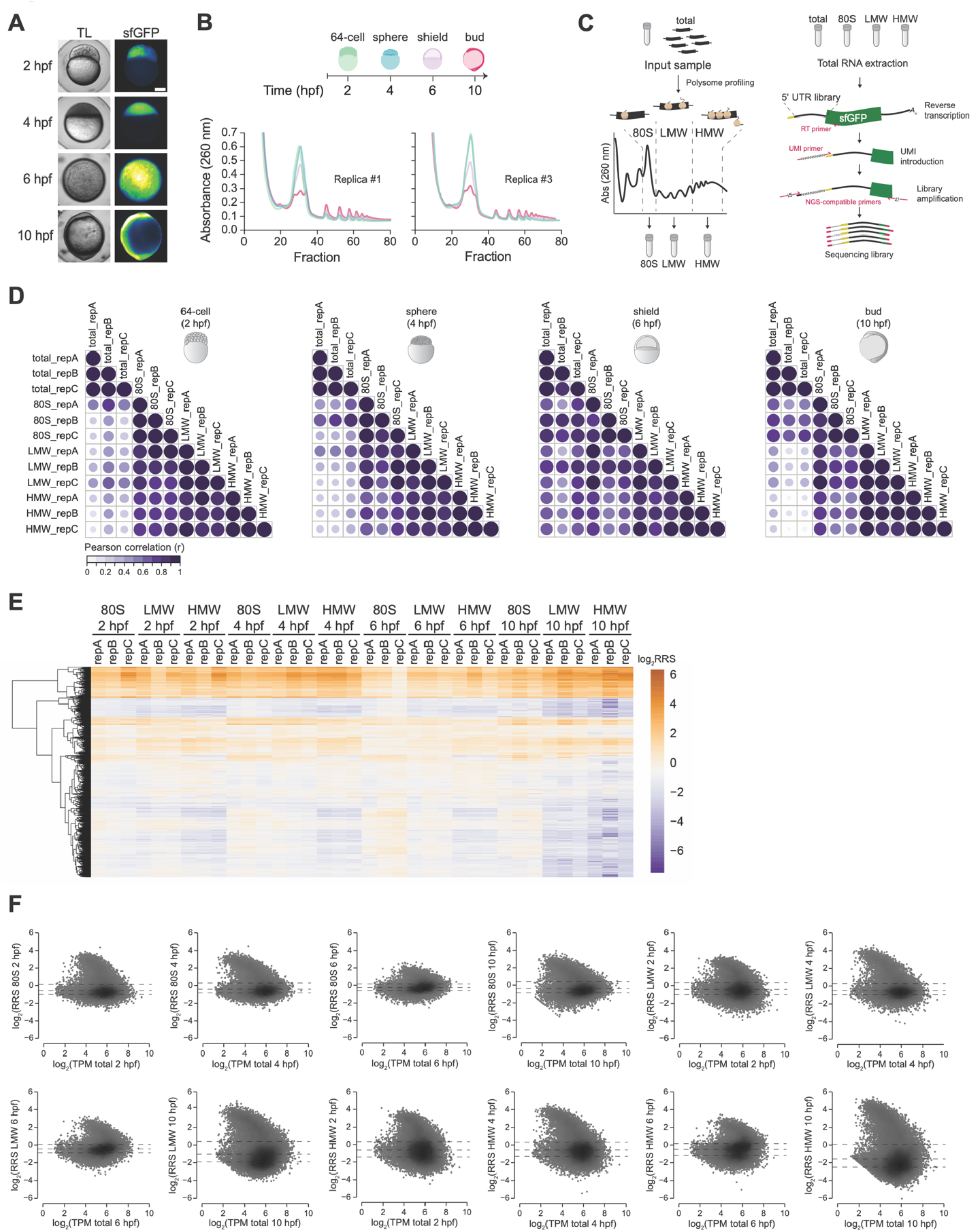
– Analysis of the 5′ UTR MPRA data, related to Figure 2. (A) Fluorescence microscopy images of representative embryos at 2, 4, 6 and 10 hpf that were injected with the pooled 5’ UTR-sfGFP mRNA library at the 1-cell stage. TL – transmitted light. (B) Polysome profiling traces for two of the three replicate experiments; replica #2 is presented in Figure 2B. (C) Schematics of the strategy for collecting input (total) and ribosome-bound RNA (80S, LMW and HMW fractions) samples, followed by total RNA extraction and generation of libraries compatible with high-throughput sequencing. (D) Pearson correlation coefficient (r) of TPM values between samples; rep = replicate. (E) Heatmap of mean log2 transformed RRS values of all replicates clustered by 5’ UTR sequence (rows) and ordered by fraction and developmental time-point (columns) (n = 17,879). (F) MA plots of mean log2 transformed TPM total values (input sample) (x axis) and mean log2 transformed RRS values (y axis) for each fraction (80S, LMW and HMW) at each time-point (2, 4, 6 and 10 hpf). See also Table S2 and STAR Methods.

**Figure S3.**
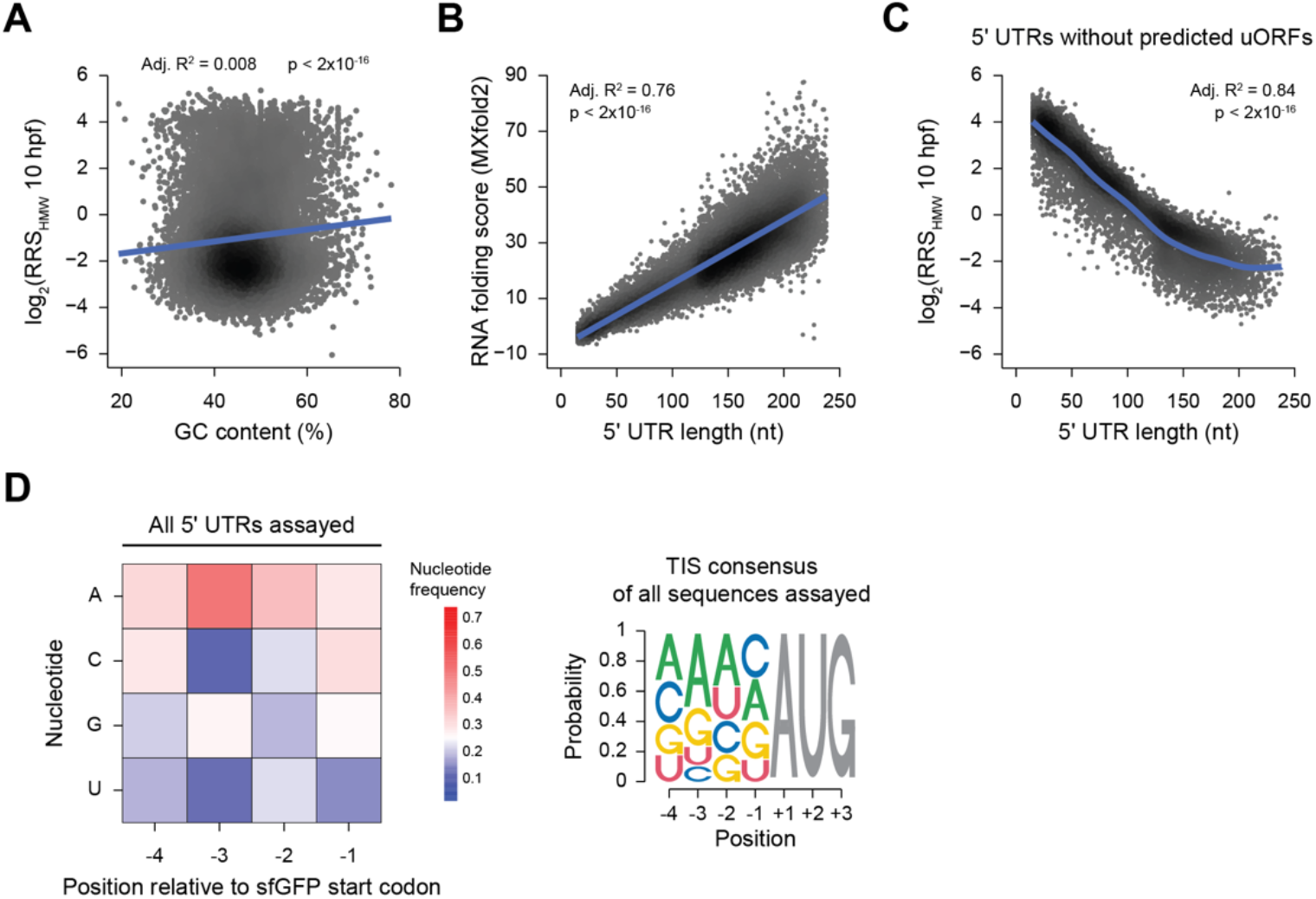
– Analysis of 5′ UTR features, related to Figure 3. (A) Effect of 5’ UTR GC content on ribosome recruitment. Scatter plot of percent GC content of the 5’ UTR sequence (x axis) and mean log2 transformed RRS values (y axis) for the HMW fraction at 10 hpf. Each dot represents one 5’ UTR (n = 17,906) and adjusted R^2^ value for best-fit to a linear model and corresponding p-value is presented. (B) Relationship between 5’ UTR length and structure. Scatter plot of 5’ UTR length (x axis) and RNA folding scores calculated by MXfold2 (y axis) for the HMW fraction at 10 hpf. (C) Effect of 5’ UTR length on ribosome recruitment is not dependent of the presence of upstream open reading frames (uORFs). Scatter plot of 5’ UTR length in nucleotides (x axis) and mean log2 transformed RRS values (y axis) for the HMW fraction at 10 hpf for 5’ UTRs without predicted uORFs. Each dot represents one 5’ UTR (n = 6,940) and adjusted R^2^ values for best-fit to a nonlinear generalized additive model and corresponding p-value are presented. (D) Mean nucleotide frequency at positions −4 to −1 of the reporter canonical sfGFP translation start site for all 5’ UTRs assayed. The position weight matrix of the translation initiation site (TIS) consensus sequence derived from mean nucleotide frequencies is presented. Grey AUG represents the translation start site of sfGFP. See also Table S3.

**Figure S4.**
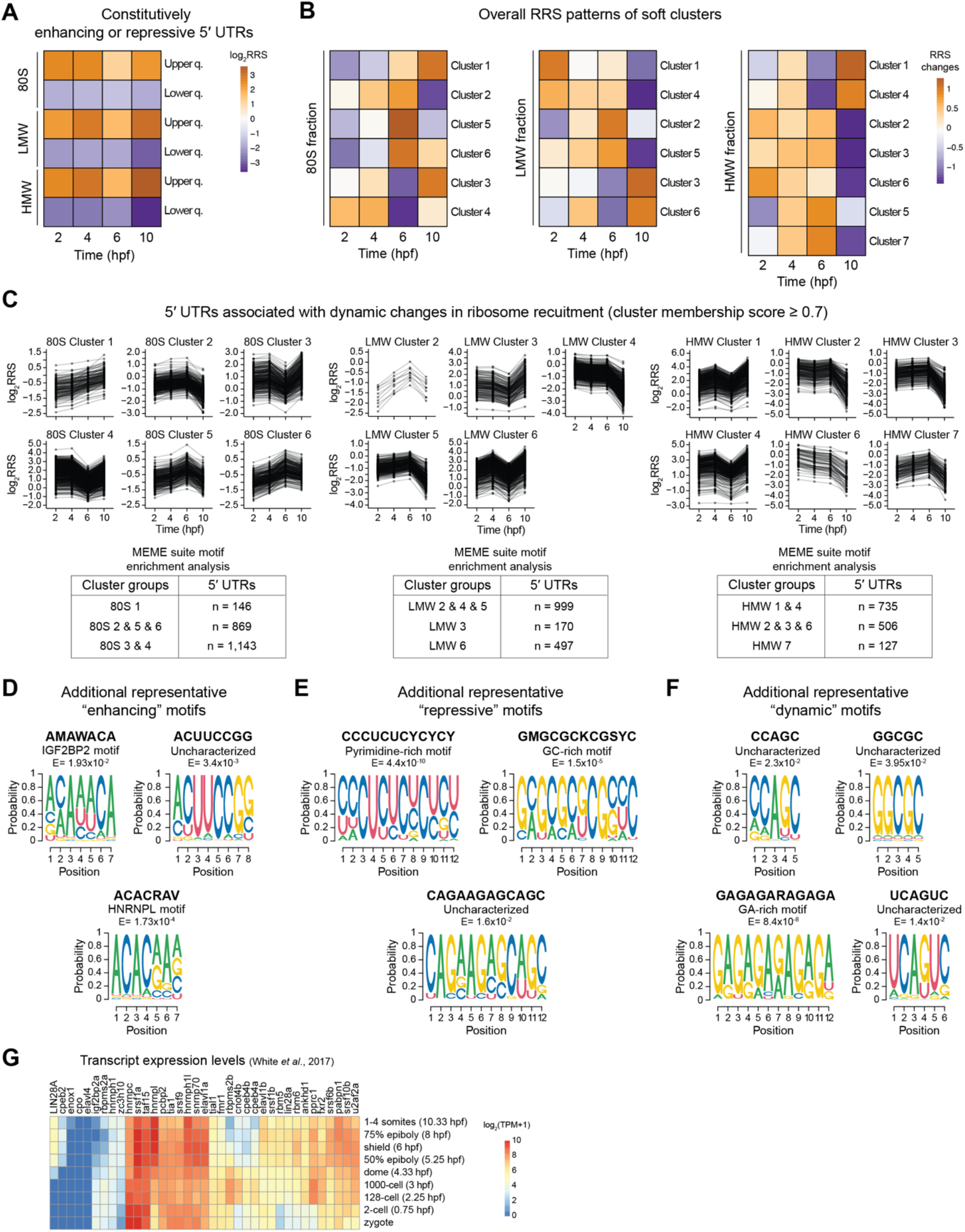
– Motif-based sequence analysis of 5′ UTR sequences, related to Figure 4. (A) Mean log2 transformed RRS values of 5’ UTRs present in the upper quantile (top 10%) or lower quantile (bottom 10%) of each fraction (80S, LMW and HMW) during the developmental time-course (2, 4, 6 and 10 hpf). (B) Soft cluster centroids across the developmental time-course. Colors represent the weighted sum of all cluster members and show the overall RRS patterns of clusters. (C) Time-series line-plots of mean log2 RRS values for each 5’ UTR in each cluster (membership score ≥ 0.7). Tables depict groups of 5’ UTR sequences used for motif enrichment analysis using the MEME suite tool (Bailey *et al*., 2015). (D) Additional representative motifs enriched in 5’ UTRs that are consistently found in the upper quantile (top 10%, “enhancing”) or (E) the lower quantile (bottom 10%, “repressive”) throughout the developmental time-course, or (F) enriched in 5’ UTRs present in the soft clusters (membership score ≥ 0.7) (“dynamic”). (G) Heatmap depicting expression values (transcript per million, TPM) (White *et al*, 2017) of transcripts encoding zebrafish RNA-binding proteins during embryogenesis that are homologous to human and *Drosophila* RNA-binding proteins with motifs matching those uncovered by motif-enrichment analysis in our study. See also STAR Methods and Table S3.

**Figure S5.**
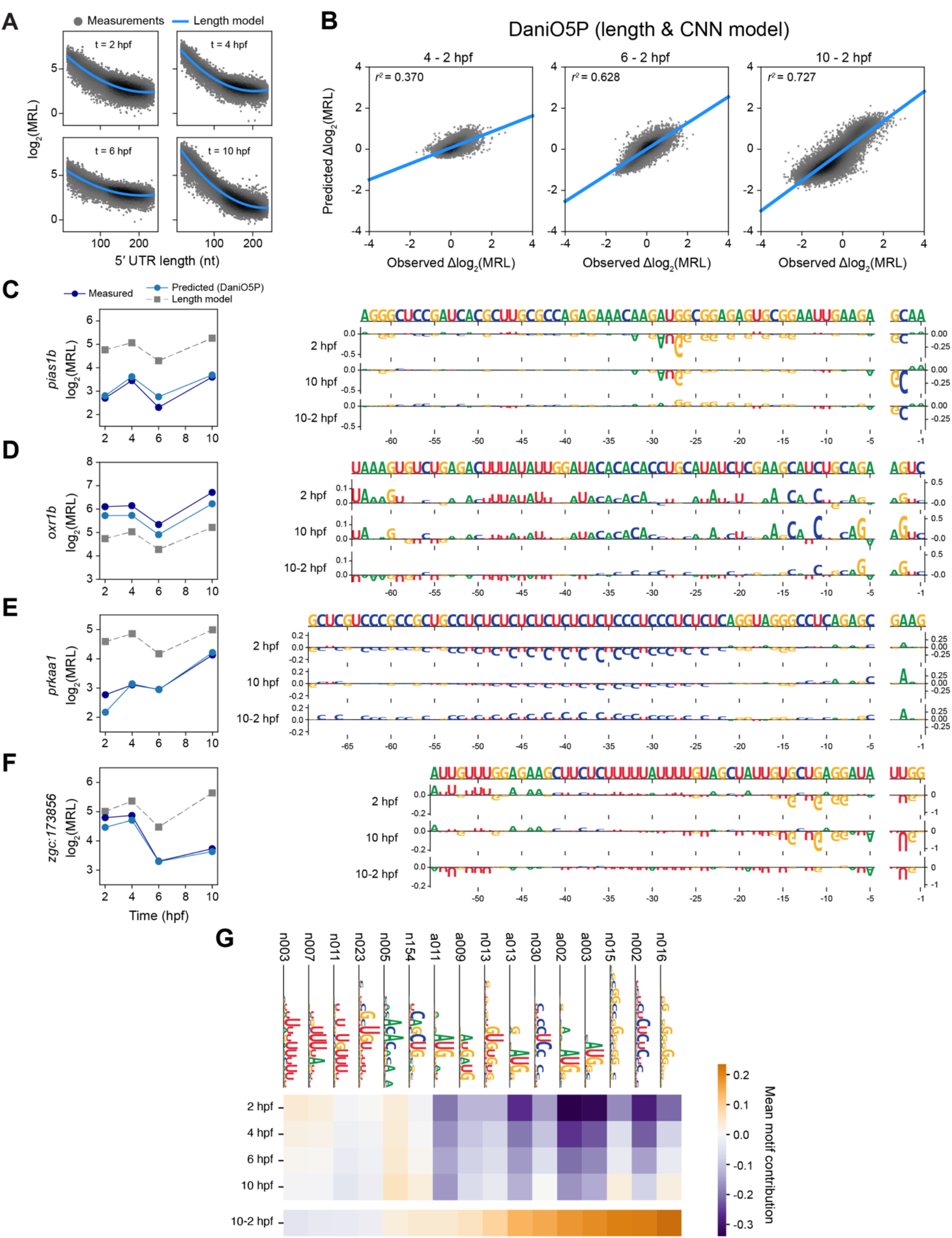
-DaniO5P model performance and additional analysis of the 5′ UTR MPRA data using deep learning, related to figure 5. (A) MRL as a function of 5’ UTR length, for all MPRA 5’ UTRs, along with second-order polynomial fits (length model). (B) Prediction performance of DaniO5P (combined length + CNN models) with respect to MRL differences between 4 and 2 hpf, 6 and 2 hpf, and 10 and 2 hpf, showing that the model captures translation dynamics especially for larger time intervals. (C-F) Additional examples of 5’ UTRs where measured and predicted MRL are substantially lower (C), higher (D), or show increasing (E) and decreasing dynamics (F) that are not predicted by length alone. Left: MRL measurements and predictions of the length model and Danio5P (length + CNN models). Right: Contribution scores (SHAP values). Note that the y axis scale was adjusted differently for the TIS context region and the rest of the sequence. (G) Motifs extracted from the CNN model ensemble, along with their mean contributions to MRL at 2, 4, 6, and 10 hpf, and to the MRL difference between 10 and 2 hpf. See also STAR Methods and Table S4.

**Figure S6.**
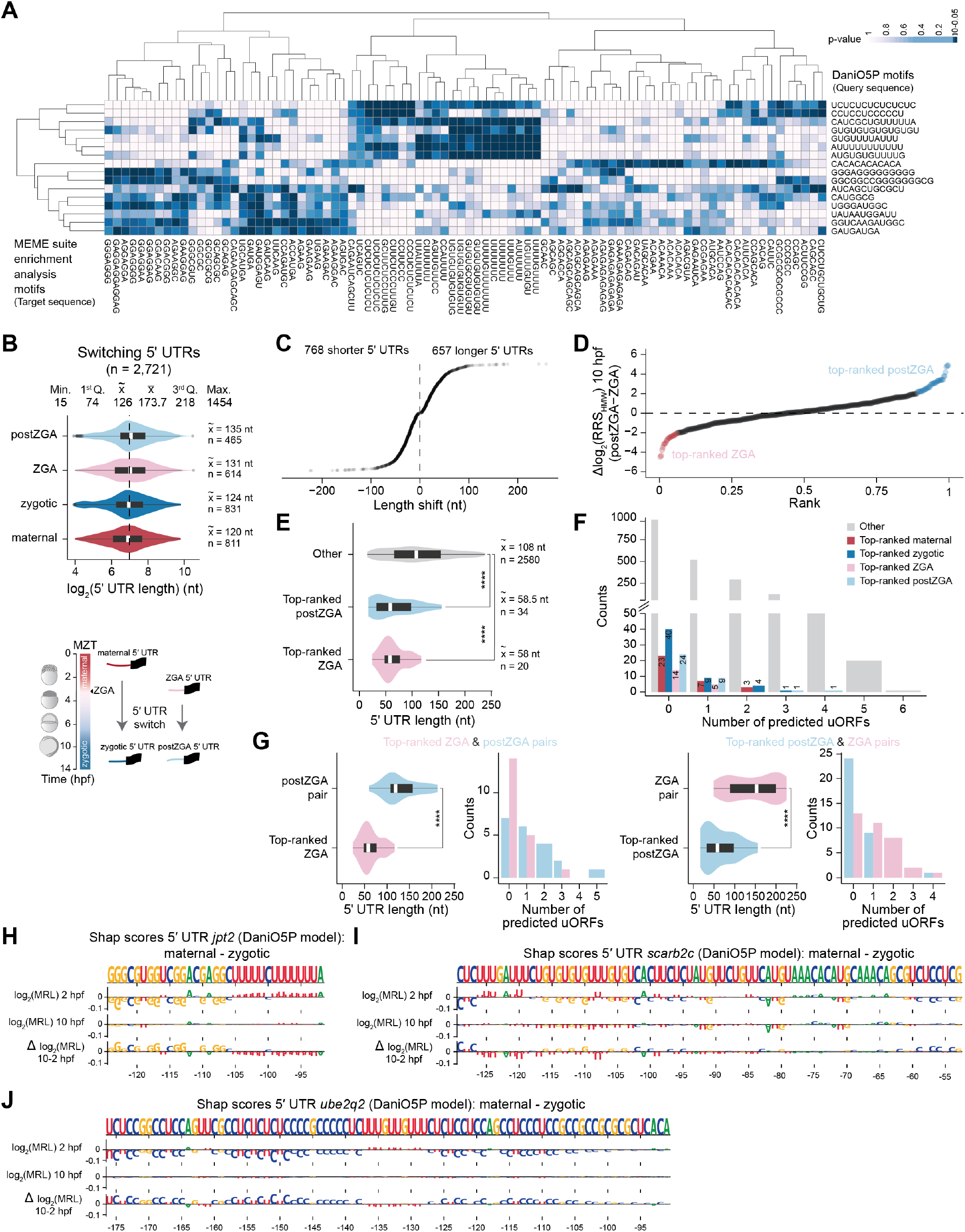
– Additional analysis of 5’ UTR isoforms in the MPRA data, related to figure 6. (A) Heatmap depicting p-values resulting from the comparison of DaniO5P model derived and MEME derived motifs using the motif comparison tool Tomtom (Gupta *et al*, 2007). (B) Length distribution of switching 5’ UTR isoforms represented in the MPRA (n = 2,721). Center line of boxplot represents median, box limits represent the upper and lower quartiles. Summary statistics of 5’ UTR lengths (in nucleotides) are provided. Switching events can take place between maternal isoforms that are deposited from the mother’s egg and re-expressed during the maternal-to-zygotic transition (MZT) as zygotic isoforms or between isoforms expressed during early phases of zygotic genome activation (ZGA) and re-expressed after ZGA (postZGA). (C) Shifts in 5’ UTR length resulting from differential transcription start site utilization throughout embryogenesis. (D) Difference in mean log2 transformed RRS values for the HMW fraction at 10 hpf between ZGA and postZGA switching 5’ UTR isoforms pairs. Cases for which the ZGA 5’ UTR isoform leads to higher ribosome recruitment (Δlog2(RRS) ≤ −2) are marked as “top-ranked ZGA” (n = 20) and for which the postZGA leads to higher ribosome recruitment (Δlog2(RRS) ≥ 2) are marked as “top-ranked postZGA” (n = 34). Only 5’ UTRs shorter than 239 nts were considered. (E) Violin plots of the length distribution of top-ranked ZGA (n = 20), top-ranked postZGA (n = 34) and other (n = 2,580) switching isoforms. Center line of boxplot represents median, box limits represent the upper and lower quartiles. **** p-value < 0.001, two-sample rank sum Wilcoxon test. (F) Bar plot depicting the number of top-ranked maternal, top-ranked zygotic, top-ranked ZGA, top-ranked postZGA and other isoforms with 0 to 6 predicted uORFs in their 5’ UTR sequences. (G) Violin plot of the length distribution and bar blot of the number of uORFs for top-ranked ZGA isoforms and respective postZGA pairs (left) and for top-ranked postZGA isoforms and respective ZGA pairs (right). Center line of boxplot represents median, box limits represent the upper and lower quartiles. **** p-value < 0.001, two-sample rank sum Wilcoxon test. (H) Nucleotide contributions (SHAP values) for the 5’ UTR variant region between maternal and zygotic isoforms at 2hpf, 10hpf, and their difference for transcripts jpt2, (I) scarb2c and (J) ube2q2. See also Table S4, S5 and STAR Methods.

**Figure S7.**
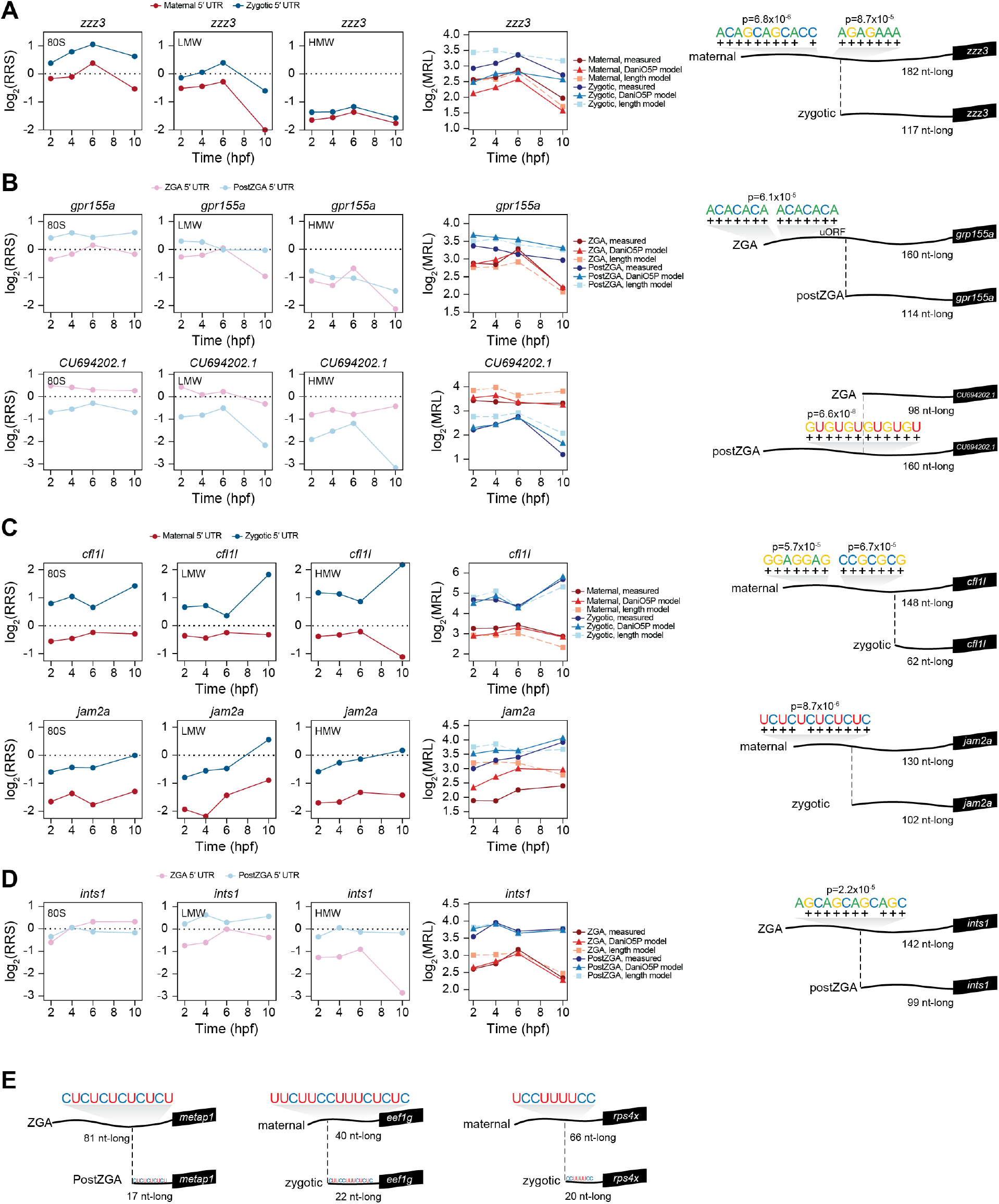
– Additional examples of switching 5’ UTR isoforms with distinct motif grammar, related to figure 6. (A-D) Left: Line plots of mean log2 transformed RRS values for 80S, LMW and HMW fractions for examples of maternal and zygotic switching 5’ UTR pairs or ZGA and postZGA switching pairs with distinct translational dynamics. Middle: Line plots of measured and predicted log2 transformed MRL values for length and DaniO5P (length + CNN) models. Right: Schematics depicting motif sequence matches and respective p-values (E) Three example transcripts for which 5’ UTR switching gives rise to a 5’ terminal oligopyrimidine (5’ TOP) motif, defined as a +1 cytidine directly adjacent to the 5’ cap of the mRNA followed by a stretch of 4-16 pyrimidines (Meyuhas & Kahan, 2015).

## STAR Methods

### Alignment and mapping of CAGE data

Raw sequenced CAGE tags (27bps) from a published zebrafish 12 developmental stages time-course (Nepal *et al*., 2013) were downloaded and mapped to a reference zebrafish genome (GRCz11, excluding alternate loci scaffolds) using Bowtie (Langmead & Salzberg, 2012) with default parameters as described in Haberle *et al.,* 2014 (Haberle *et al*., 2014). All analysis were conducted in the R statistical computing environment (R Core Team, 2018; https://www.R-project.org/) using Bioconductor (Gentleman *et al*, 2004); http://www.bioconductor.org) software packages.

### CTSS calling

CAGE tag-defined transcription start site (CTSS) calling and positional frequency determination was performed using the CAGEr software package (Haberle *et al*., 2015). To enable comparison between multiple samples, raw tag count was normalized to a reference power-law distribution (Balwierz *et al*, 2009) based on a total of 10^6^ tags and *α* = 1.2, resulting in normalized tags per million. Low fidelity TSSs supported by less than 2 normalized tag counts were filtered out. Individual TSSs were clustered into tag clusters (TCs) using the data-driven parametric clustering method paraclu (Frith *et al*, 2008) (clusters merged to non-overlapping), and clusters supported by a normalized signal > 5 tpm in at least one developmental stage were included. TCs for which the ration between the maximal and minimal density was lower than 2 (i.e. stability < 2) and longer than 200 bps were excluded. TCs across all developmental stages within 100 bp of each other were aggregated into a single promoter region.

### Recovery of 5′ UTR sequences

Gene annotations corresponding to the zebrafish genome build GRCz11 were retrieved from the Ensembl database (Ensembl 100 release) (Kinsella *et al*, 2011). CAGE-recovered TCs for which the dominant TSS is more than 500 bp away from the closest annotated TSS were excluded. To retrieve precise and stage specific TSS data, we considered the dominant (most frequently used) TSS positions for each of the 12 developmental stages as the 1^st^ nucleotide position of the 5′ UTR. The nucleotide position immediately upstream of the ATG triplet of annotated coding sequence (CDS) start sites was considered the last nucleotide position of the 5′ UTR sequence. Coding transcripts lacking an annotated CDS start site or for which the dominant TSS falls downstream of the annotated CDS start site were excluded. Similarly, transcripts for which the dominant TSS falls within an intron were not considered for the analysis, since we could not confidently determine the downstream used splice acceptor site. We assume these correspond to expressed splice variants. To identify differential TSS usage across samples, we scored “shifting promoters” (shifting score threshold = 0.6, K-S test, FDR :: 0.01) using the CAGEr package (Haberle *et al*., 2015). 5′ UTR sequences were extracted using BEDTools (Quinlan & Hall, 2010).

### 5′UTR library design and synthesis

The distance between the exit channel of the ribosome and the P site is ∼12 nts, thus a 5′ UTR length of <15 nts is expected to produce a 48S PIC complex in which the m^7^G cap will be situated in the mRNA exit channel and hamper engagement of the exit channel with nucleotides 5′ of the AUG, impairing their translation (Hinnebusch, 2011). As such, recovered 5′ UTRs shorter than 15 nts were filtered out. For transcripts undergoing switches in TSS usage, we included all switching unique 5′ UTR variants (n = 2,721). For transcripts with more than one dominant TSS within a promoter region but that did not score as switching, we considered the longest 5′ UTR isoform (n = 11,620). After filtering steps, the UTR library represented 10,354 genes and 11,445 transcripts. For 5′ UTRs :: 238 nt long (n = 9,863), the full “endogenous” sequence was considered. For 5′ UTRs > 238 nt long (3,446), sequences were split into equally long stretches flanked by 10 nt of flanking overlapping regions (Figure S1C). After splitting, the 5′ UTRs consisted of a total of 18,154 unique sequences ranging from 15 to 238 nt in length. For library amplification and cloning purposes, common upstream (PBS1, GAAGAGTAGCCTGCAGATAGAC, 22 nts) and downstream (PBS2, ATGGTGTCTAAAGGAGAGGAGC, 22 nts) sequences were introduced to flank each 5′ UTR and a third common sequence (PBS0, TGGTTGATTACGGTCGCA, 18 nt) was introduced at the most 3′ UTR of the sequence (Figure S1D). In cases where the original 5′ UTR was shorter that 238 nt in length, a random sequence (“filler”, in pink color) was added between PBS2 and PBS0 for a total length of 300 nts. The 300-nt-long pooled oligonucleotide library (n = 18,154) was synthesized by Twist Bioscience.

### 5′ UTR library cloning and mRNA synthesis

The library cloning strategy was adapted from the STARR-seq cloning protocol (Neumayr *et al*, 2019). A pMB1vector (Twist Bioscience) custom construct was designed to include an SP6 promoter followed by a landing site for directional cloning of the 5′ UTR library, the sfGFP open reading frame and a 3′ UTR zebrafish sequence (BG1, 110 nt long) devoid of known regulatory sequence motifs (Rabani *et al*., 2017) (pMB1-BG1 vector) (Figure S1D and E). The synthetic oligonucleotide pool (Twist Bioscience) was amplified via the two primer handles (PBS0 and PBS1, 14 cycles) to yield a 300 bp long PCR product, followed by a low-cycle nested PCR (PBS1 and PBS2, 7 cycles) to generate the 5′ UTR library ranging from 15 (59 bp PCR product) to 238 nt-long (282 bp PCR product) sequences using the KAPA HiFi HotStart ReadyMix (Roche). The PCR product (5′ UTR insert pool) was purified using Angecourt AMpure XP beads (1.8 bead volume to PCR reaction volume) with the addition of isopropanol to 45%. The pMB1-BG1 vector was digested with BbsI-HF (NEB) to remove a filler sequence flanked by inverted BbsI restriction sequences and to allow for directional cloning. The digested vector was purified with the QIAquick PCR purification kit (QIAGEN) and purified again with the MinElute PCR purification (QIAGEN). The 5′ UTR insert pool was cloned into the BbsI-digested pMB1-BG1 vector between the SP6 sequence and the sfGFP ORF, to restore the 5′ UTR of the sfGFP. Homologous recombination directional cloning was performed using In-Fusion HD (Takara Bio). The cloned DNA library was electroporated into electrocompetent MegaX DH10B bacteria (Invitrogen) at a 100x library coverage. The colony lawn resulting from transformation was harvested and plasmids extracted using the Plasmid Plus Giga kit (QIAGEN). The plasmid library was PCR amplified using a SP6 forward primer (Fwd_IVT_SP6: 5′ CACGCATCTGGAATAAGGAAGTGC 3′) and a 3′ UTR-specific reverse primer containing a 36 nt-long T overhang (Rev_IVT_BG1orig: 5′ TTTTTTTTTTTTTTTTTTTTTTTTTTTTTTTTTTTTTCCTGTGAGTCCCATGGGTTTAAG 3′), followed by DpnI treatment (1h 37°C) and PCR product purification using Angecourt AMpure XP beads (0.8 bead volume to PCR reaction volume) (Figures S1D and S1E). The PCR product encompassed 99.9% (18,142/18,154) of the designed 5′ UTR sequences with lengths ranging from 15 to 238 nts (Figures S1F and S1G). This PCR template was used for *in vitro* transcription (mMessage Machine Sp6 kit, Thermo Fisher) by incubation on a thermocycler machine for 12h at 30°C. After incubation, DNase treatment was performed according to kit instructions (mMessage Machine Sp6 kit, Thermo Fisher) and the IVT mRNA product was purified using the RNA Clean & Concentrator kit (Zymo Research). The resulting IVT product is a 5′ capped reporter mRNA pool consisting of variant 5′ UTRs driving translation of an invariant sfGFP ORF, the 3′ UTR and a 36 nt-long polyA tail (Figure S1H). The mRNA library faithfully represents 5′ UTR sequences and consists of almost the entirety of sequences synthesized. Transcript abundances in the *in vitro* transcribed mRNA library range from ∼1 to ∼400 TPM (25^th^ percentile = 23 TPM; 50^th^ percentile = 44 TPM; 75^th^ percentile = 76 TPM) with shorter 5′ UTRs less well represented (Table S1), mirroring the less efficient cloning of shorter oligonucleotide sequences.

### Zebrafish husbandry and embryo rearing

Wild-type zebrafish embryos (*Danio rerio*, TLAB strain) were grown in standard conditions (28°C at a 14/10 hour light/dark cycle). The TLAB strain was generated by crossing zebrafish natural variant TL (Tupfel Longfin) with AB strain. Zebrafish embryos were collected from TLAB strain crosses, incubated in E3 medium (5 mM NaCl, 0.17 mM KCl, 0.33 mM CaCl2, 0.33 mM MgSO4, pH 7.2) in standard conditions (28°C at a 14/10 hour light/dark cycle) and staged as described (Kimmel *et al*, 1995). All experiments were conducted according to federal guidelines for animal research and approved by the Kantonales Veterinäramt of Kanton Basel-Stadt (under the Animal Holding License Form 1035H).

### Image acquisition and quantification

Embryos were injected at the 1-cell stage and allowed to develop in standard conditions. For 5′ UTR mRNA library pool experiments, 80 pg/embryo were injected. For single-reporter injections, 80 pg/embryo were injected (40pg 5′ UTR-sfGFP test reporter co-injected with 40pg mCherry control reporter). Embryos were collected at the desired developmental stage and placed on a custom-made agarose mold with squared indents for placing and aligning the embryos. For fluorescence intensity quantification, zebrafish embryo images were acquired using an upright ZEISS Axiozoom coupled to an Axiocam 503 color/mono digital camera (14-bit depth) in black & white color mode with fixed laser power, fixed zoom and fixed exposure time for red mRF12 (590/612) and green AF488 (493/517) channels. Images were quantified using Fiji (Image J) by selecting the region of the embryo’s cells with mCherry signal and measuring mean pixel intensity for the red channel, and applying the same region of interest area for measuring mean pixel intensity for the green channel. Background signal for each channel was subtracted from the mean pixel intensity for each image. Normalized fluorescence intensity values consist of the ratio between corrected mean pixel intensities of green and read channels (sfGFP/mCherry). Two rounds of single-reporter injections (embryos from two different clutches) were performed for fluorescence intensity quantification.

### Zebrafish embryo injections and polysome profiling

Embryos were injected at the 1-cell stage with 80 pg of the 5′ UTR mRNA library pool and allowed to develop in standard conditions. Embryos were staged and collected at the 64-cell (2 hpf), sphere (4 hpf), shied (6 hpf) and bud (10 hpf) stage. For each replicate sample, staged embryos were pooled and flash-frozen in liquid nitrogen and stored at −80°C. Three independent time-courses were performed (1 time-course per replicate, on three different days). For each polysome profiling experiment, 100 embryos injected with 5′ UTR mRNA library were used. Immediately before polysome gradient preparation, embryos were lysed in 450 μL polysome gradient buffer (20 mM Tris-Cl pH 7.5, 30 mM MgCl2, 100 mM NaCl, 0.25 % Igepal-630 [v/v], 0.1% sodium deoxycholate (w/v), 100 μg/mL cycloheximide, 0.5 mM dithiothreitol [DTT], 1 mg/mL heparin and 40U/mL recombinant RNasin ribonuclease inhibitor [Promega]) with ∼30 strokes of a motor-driven “B” pestle while on ice. After lysis, 7.2U of TURBO DNase (Invitrogen) were added and the embryo lysate was incubated for 15 min at 4°C on a rotating mixer. Samples were centrifuged at 13,000 rpm for 10 min at 4°C, and 400 μL of clarified lysate were loaded onto a 5-50% (w/v) sucrose gradient prepared in TMS buffer (20 mM Tris-Cl pH 7.5, 5 mM MgCl2, 140 mM NaCl, 100 μg/mL cycloheximide, 1 mM DTT). The remaining 50 μL of embryo lysate were kept as input sample. Gradients were centrifuged in a SW40 Ti rotor (Beckman) at 4°C for 150 min and profiles were analyzed using a Gradient Station (Biocomp Instruments) with continuous recording of optical density (OD) at 260 nm.

### Polysome fraction processing

Fractions of 1200 μL were individually collected and SDS was immediately added to a final concentration of 1%. Fractions corresponding to the 80S peak (fractions 4 + 5), to low-molecular weight (fractions 6 + 7) or high-molecular weight (fractions 8 + 9) polysomes were pooled together in 15mL RNase-free tubes (Ambion) (Figure S2C). A volume of 2.4 mL TRI Reagent (Sigma) was added per tube, vortexed and stored at −80°C. To the withheld input sample (50 μL), 1 mL TRI Reagent (Sigma) was added, vortexed and stored at −80°C. For total RNA extraction, samples were thawed at room temperature. For fractionated samples, half of the sample (∼2.4 mL) was transferred to a fresh RNase free 5mL Eppendorf tube and an extra 1.25 mL of TRI Reagent (Sigma) was added to each, for a final ratio of ∼2:1 TRI Reagent:sample. Samples were mixed by vortexing and incubated at room temperature for 5 min. To each 5 mL tube, 500 μL of chloroform (without isoamyl alcohol or other additives) were added (or 200 μL to the input sample) and samples were vortexed. Samples were centrifuged at 12,000 g (an Eppendorf 5427 R centrifuge with a rotor for 5 mL tubes) for 25 min at 4°C. The aqueous phase was transferred to a fresh 5 mL Eppendorf tube and an equal volume of pure ethanol (> 99%) was added followed by vortexing. The Direct-zol RNA Miniprep kit (Zymo Research) was used for RNA purification. The mixture was transferred into a Zymo-Spin IICR Column in a collection tube and centrifuged for 30 sec at 15,000 rpm. The procedure was repeated until all sample was passed through the column (∼ 850 μL at a time). The Direct-zol RNA Miniprep kit protocol was followed, excluding the DNase treatment step. The final sample was eluted in 25 μL DNase/RNase-free water, 2 μL of sample were collected for gel quality control and the remaining was stored at −80°C. One sample of total RNA from non-injected embryos was included as a negative control for the procedure described below.

### Library preparation and high-throughput sequencing

A total of 50 sequencing libraries were prepared: replica A (input, fractions 4 + 5, 6+7 and 8+9) at 2, 4, 6 and 10 hpf, replica B (input, fractions 4 + 5, 6+7 and 8+9) at 2, 4, 6 and 10 hpf, replica C (input, fractions 4 + 5, 6+7 and 8+9) at 2, 4, 6 and 10 hpf, as well as libraries of the pooled PCR template used for library *in vitro transcription* and the uninjected 5′ UTR mRNA library to determine library complexity. For samples corresponding to replicas A-C and the uninjected 5′ UTR mRNA library, an RT reaction containing 11 μL of RNA sample and 1 μL transcript specific primer (2 μM RT_TSP primer: 5′ GTTCAGACGTGTGCTCTTCCGATCTGAACTTGTGGCCGTTCACGTCTCCAT 3′) targeting the beginning of the sfGFP open reading frame was performed (55°C for 1h, then 70°C for 15 min and hold at 4°C) using SuperScript II Reverse Transcriptase (Thermo). This was followed by an RNase treatment to remove template mRNA by adding 0.2 μL RNase A (10 mg/ml; Thermo Scientific EN0531) per 20 μL RT reaction, and incubation for 1h at 37°C. The cDNA was purified with AMPure XP beads (1.8 vol beads to 1 vol cDNA volume) and eluted in 22 μL DNase/RNase-free water. Then, 20 μL pure cDNA sample were used for UMI introduction by linear PCR using KAPA HiFi HotStart ReadyMix (Roche) at the second strand synthesis step (P5_UMI_fwd: 5′ CACGACGCTCTTCCGATCTNNNNNNNNNNGAAGAGTAGCCTGCAGATAGA*C 3′, where * denotes a phosphorothioate bond), analogous to the procedure described in Neumayr *et al*., 2019. The AMPure XP beads purification step was repeated (0.8 vol beads to 1 vol PCR product) to remove excess primer and eluted in 22 μL DNase/RNase-free water. Library amplification was performed with primers containing TruSeq indexes comptible with Illumina high-throughput sequencing (Illumina i5: 5′ AATGATACGGCGACCACCGAGATCTACAC-i5-ACACTCTTTCCCTACACGACGCTCTTCCGATCT 3′; Illumina i7: 5′ CAAGCAGAAGACGGCATACGAGAT-i7-GTGACTGGAGTTCAGACGTGTGCTCTTCCGATCT 3′). First, qPCR reactions using a fraction of the UMI-tagged PCR product as input (1.25 μL in 25 μL qPCR reaction) were performed to determine the ideal number of cycles used for amplification of each library. The qPCR reactions were ran using The KAPA HiFi HotStart ReadyMix (Roche) with EvaGreen Dye (Biotium) and ran in parallel for all samples up to 40 cycles, and the cycle number at which the ampification reached its exponential phase for each sample was noted (CN_test_).

The final library amplification was performed using 10 μL UMI-tagged PCR product and the KAPA HiFi HotStart ReadyMix (Roche), using the same cycling conditions and a total number of cycles corresponding to CN_test_ - 3 tailored to each sample. The PCR product (ranging between 263-486 bp in length) was purified with AMPure XP beads (0.8 vol beads to 1 vol PCR product) and 5 μL were analysed by gel electrophoresis. For generating a library of the pooled PCR template (template for *in vitro* transcription of the 5′ UTR mRNA library), a PCR reaction targeting the variant 5′ UTR sequence of the 5′ UTR library plasmid DNA pool (Fwd_IVT_SP6: 5′ CACGCATCTGGAATAAGGAAGTGC 3′ and NGS_TSP_rev: 5′GTTCAGACGTGTGCTCTTCCGATCTGAACTTGTGGCCGTTCACGT3′) was performed.

The PCR product was treated with DpnI and purified with AMPure XP beads (0.8 vol beads to 1 vol PCR product), followed by UMI introduction by linear PCR and library preparation as described above. Library quality control and quantification was performed using a 5200 Fragment Analyzer System (Agilent). The 50 libraries were sequenced with 100 nt paired-end reads (PE100) with uneven loading for the forward read in a NovaSeq machine. Library quality control and high-throughput sequencing was performed by the Genomics Facility Basel (D-BSSE/ETHZ).

### Sequence processing, TPM and RRS calculations

Sequencing data was filtered to retain only sequences with a UMI immediately followed by the common adaptor sequence (GAAGAGTAGCCTGCAGATAGAC, colored in yellow in Figure S1D) (see also Table S1, sheet *sequencing_stats*). UMIs were extracted using UMI-tools (Smith *et al*, 2017). Adaptors (invariant common adaptor and the sfGFP sequences flanking the variant 5′ sequence) were partially trimmed using the non-internal adaptor trimming method of Cutadapt (Martin, 2011), , leaving each 5′ UTR sequence flanked by 10 invariant bps. Leaving parts of the invariant upstream and downstream flanking sequences allows to map 5′ UTR isoforms that differ in size but share much of their sequence. Up to 2 mismatches were allowed for the remaining common adaptor sequence and 3 for the remaining sfGFP sequence. Bowtie2 (Langmead & Salzberg, 2012) was used to align retained reads to a reference set containing all 5′ UTRs in the synthetic oligonucleotide library (18,154 sequences) flanked by the common adaptor sequence (upstream) and sfGFP open reading frame sequence (downstream) using the “end-to-end” and “very-sensitive” parameters. Multi-mapped reads were removed by applying a MAPQ>=2 and flag 256 filter using SAM-tools (Li *et al*, 2009). Reads were deduplicated using UMItools (Smith *et al*., 2017) and the number of reads mapped to each 5′ UTR entry were recorded. For determining TPM values, the number of mapped reads for each 5′ UTR entry were divided by the total number of mapped reads in the library and multiplied by 10^6^ (+ one pseudo count). For determining RRS values, TPM values for each 5′ UTR entry in the library fraction sample were divided by the TPM values of the respective 5′ UTR entry in the input sample (see Figure 2C and S2C), in a manner analogous to that described in (Niederer *et al*., 2022). A TPM ≥ 2 cutoff was applied for RRS calculations. RRS values were log_2_ transformed, and the arithmetic mean of log_2_RRS for each 5′ UTR entry of all three replicate samples was calculated. Entries for which at least one of the replicate values was NA were excluded.

### RNA structure and uORF predictions

RNA folding scores for each assayed 5′ UTR sequence were predicted by inputting 5′ UTR FASTA sequences into MXfold2 (https://github.com/mxfold/mxfold2, version mxfold2-0.1.1) employing the default parameters trained from TrainSetA and TrainSetB (Sato *et al*., 2021). The number of predicted uORFs for each assayed 5′ UTR sequence was determined with the R Package ORFik (https://bioconductor.org/packages/release/bioc/html/ORFik.html) (Tjeldnes *et al*., 2021). Upstream ORFs consisting of a uAUG and a downstream in-frame stop codon were searched in the 5′ to 3′ direction in all three possible reading frames using the “minimumLength=0” filter (START+STOP = 6 bp minimum ORF length).

### Data subsetting for 5′ UTR sequence analysis

Data subsetting was performed by quantile analysis and by fuzzy-c-means clustering (Figure 4A). Deciles of log_2_RRS values for each fraction (80S, LMW and HMW) at each developmental time-point (2, 4, 6, and 10 hpf) were determined. 5′ UTR entries that were consistently found either in the top 10% or bottom 10% deciles across all time-points for either the 80S, LMW or HMW fractions were filtered and used for motif-based sequence analysis (see Table S3). Fuzzy-c-means clustering was performed using the Mfuzz package (Kumar & M, 2007), using the weighted k-nearest neighbor method to replace missing values and setting the fuzzifier clustering parameter m to 2.502999. Values were filtered *a posteriori* by setting a membership score threshold of α ≥ 0.7. 5′ UTR entries displaying analogous changes in RRS across developmental time were grouped together for motif-based sequence analysis (see Figure S4C).

### Motif-based sequence analysis

FASTA sequences of 5′ UTR entries in the upper quantiles, lower quantiles and fuzzy c-means clusters were generated. The XSTREME (Grant & Bailey, 2021) motif discovery and enrichment analysis (version 5.5.4) data submission form from the MEME suite online tool (Bailey *et al*., 2015) was used for extracting motifs (accessible via https://meme-suite.org/). Sequences obtained by quantile and clustering analysis were used for motif search, and sequences in the MPRA library (n = 18,154) were used as control sequences. XSTREME E-value threshold was set ≤ 0.05, motif width was set from 5 to 12 nts, the model of control sequences was used as background model and sequences were aligned to their right ends. STREME and MEME E-values were set to default. Output motifs and respective E-values are in Table S4 (sheet *Motif_enrichment_analysis*). For searching for identified motifs (Table S4) in 5′ UTR switching variants, motifs were uploaded onto MAST (Bailey & Gribskov, 1998) and searched using sequence E-value ≤ 10 threshold, removing motifs that are too similar to others and sorting motifs by best combined matches. In parallel, FIMO (Bailey *et al*., 2015) was used for individual motif scanning using a match p-value < 1×10^−4^ threshold. P-values of motifs matched presented in Figures 6 and S7 correspond to individual motif p-values. Motif matches are in Table S4 (sheets *MAST_motif_matches* and *FIMO_motif_matches*). For MEME-derived and CNN-derived motif comparison (Figure S6A; Table S4 sheet *Tomtom_comparison*), Tomtom (Gupta *et al*., 2007) was ran using the Pearson correlation coefficient function and a E-value < 1000 threshold.

### MRL calculation

The mean ribosome load (MRL) is meant to represent the ribosome loading of a transcript by a single number. MRL at each time-point was calculated according to the equation in Figure 5A, where n_80S_, n_LMW_, and n_HMW_ are estimated mean number of ribosomes in the 80S, LMW, and HMW fractions, respectively. In contrast to the original MRL calculation (Sample *et al*., 2019), our definition uses the total RNA TPM in the denominator instead of the unweighted TPM sum across fractions.

### DaniO5P length model

We fit second order polynomials of the form a_2_x^2^ + a_1_x + a_0_, where x is the UTR length, to each one of the following 7 quantities: log_2_(MRL) at 2, 4, 6, and 10 hpf, and Δlog_2_(TPM_total RNA_) at 4 minus 2 hpf, 6 minus 2 hpf, and 10 minus 2 hpf. Polynomial parameters are shown in Table S4.

### DaniO5P CNN model

We trained 10 convolutional neural network (CNN) models to predict, given an arbitrary input 5’UTR sequence, the residuals from the length model (measurement – length model prediction) for all four log_2_(MRL) and three Δlog_2_(TPM_total_RNA_) outputs. Input sequences were one-hot encoded with a maximum length of 238 nt, with shorter sequences padded to the left with zeros. The architecture of each CNN was a small VGG-16 (Simonyan & Zisserman, 2014) input sequences are passed through a series of convolutional blocks, each with two convolutional and one max pooling layer, followed by a fully connected layer and a linear layer with one output for each prediction. Hyperparameters were optimized individually based on performance on the validation set of chromosomal split 0 (see below). Final hyperparameter values were: number of convolutional blocks: 3, convolutional filter size: 7, number of filters in the first convolutional layer: 128, dropout in the convolutional layers: 0.1, number of units in the fully connected layer: 150, dropout in the fully connected layer: 0, all activations were ReLu. Model training was performed in python 3.9 using tensorflow 2.4 with the Adam optimizer, a mean square error penalty, and a decayed learning rate scheduler. Only sequences with a minimum TPM_total RNA_ of 2 on all replicates at all time-points were used for CNN training (17,879 total).

To generate an ensemble of diverse models and make sure the generalization ability of each individual model is accurately assessed, a cross-fold training strategy based on chromosome splits was used: For each model, sequences from two or three chromosomes were held out from training (test set, 1.8k sequences on average), two or three more were used for early stopping (validation set, 1.8k sequences on average) and the remaining ones were used for training (14.3k sequences on average). This split was designed such that each chromosome was part of the test set of exactly one model, while trying to maintain the number of sequences in the training, validation, and test sets similar across models. Designing such splits is a version of the “multiway number partition problem”, for which we used the prtpy python package (https://github.com/coin-or/prtpy). Table S4 lists the chromosomes and number of sequences in the training, validation, and test set of each model. To avoid overfitting during training, performance on the validation set was evaluated after each epoch, and training was stopped when this performance failed to improve for 10 epochs. To evaluate model performance (Figures 5C and S5B), each individual model was used to generate predictions on its own test set. For all other calculations in this manuscript, “model output” refers to the average across all 10 ensemble models.

### Calculation of nucleotide contribution scores

We obtained contribution scores with respect to all 7 model outputs of all nucleotides within all 17,879 MPRA sequences used for model training and evaluation. To this end, we used a custom version of DeepSHAP (https://github.com/kundajelab/shap; commit 29d2ffab405619340419fc848de6b53e2ef0f00c), which can generate hypothetical contribution scores needed for motif discovery via TFModisco (Shrikumar *et al*, 2018) though this was not the motif discovery method we ultimately used (see below). As a background, we calculated the dinucleotide frequencies across all MPRA sequences and precomputed 25 random sequences for each length between 15 and 238. Computation of contribution scores was performed in python 3.6 using tensorflow 1.15. Nucleotide contributions for differences in outputs (e.g. log_2_(MRL_10hpf) – log_2_(MRL_2hpf)) were obtained by subtracting the corresponding output contributions.

### CNN motif discovery

Motif discovery was performed in several steps. 1) Convolutional filter motif extraction: for each of the 128 convolutional filters in the second convolutional layer (first convolutional block, before the max pooling layer), we accumulated the 100 sequence fragments (seqlets) that resulted in maximal filter activation across all MPRA sequences, and used them to generate a filter position weight matrix (PWM). Seqlet length was set to 13, the receptive field after two convolutions with filter size 7. We required seqlets to be entirely contained within a sequence, therefore this analysis may not effectively find motifs influential at the very beginning or end of the 5′ UTR. This process resulted in 1280 filter motifs (128 motifs per model x 10 models). 2) Motif clustering: to account for motif redundancy within and across models, PWMs were clustered using a standalone version of RSAT matrix clustering (Castro-Mondragon *et al*, 2017) (https://github.com/jaimicore/matrix-clustering_stand-alone), modified by us to remove the ability to use PWM reverse complements. To improve clustering results, we first separated motifs that contained AUGs from the rest (AUG likelihood threshold = 0.83333) and clustered these sets separately with different normalized correlation thresholds (AUG motifs: 0.55, non-AUG motifs: 0.6) but otherwise identical parameters (correlation threshold = 0, minimum number of aligned positions = 4, linkage method = average). This resulted in 25 AUG and 573 non-AUG motif clusters. 3) Motif filtering and post-processing: Reproducible and robust motifs should emerge independently in models trained on different data splits. Thus, motif clusters were required to contain motif filters originating from multiple individual models (10 for AUG motifs, 5 or more for non-AUG motifs), and those that did not were discarded. This resulted in 5 and 11 AUG and non-AUG motifs, respectively. Finally, motif cluster PWMs were trimmed at one base from the first position with information content greater than 0.3 on both sides to produce the final list of motifs. 4) Motif contribution score calculation: Motif cluster PWMs were scanned against all MPRA sequences with contribution scores using FIMO with the following options: pvalue < 1e-4, given strand only, background model from motif file, don’t parse genomic coordinates. For each match in each motif, contribution scores were summed across all matching bases and averaged across all matches to obtain the motif contribution scores. The final motifs and their contributions are shown in Figure S5G.

### Statistical analyses and plotting

Statistical analyses relative to plots presented in Figures 3, 6C-E, S2D, S3, S6E and S6G were performed in R version 4.3.1 (R Core Team, 2021). Statistical analyses relative to plots presented in Figure 2H were performed in GraphPad Prism. Hierarchical clustering (Figures 2D and S2E) was performed with the R package pheatmap (https://cran.r-project.org/web/packages/pheatmap/pheatmap.pdf), Fuzzy c-means clustering was performed with the R package Mfuzz (Kumar & M, 2007) and motif logos were generated with the R package SeqLogo (Bembom & Ivanek, 2023). Statistical analyses relative to CAGE data were performed using the R package CAGEr (Haberle *et al*., 2015). Motif p-values, E-values and PWMs were outputted from MEME suite tool (Bailey *et al*., 2015) analyses. For CNN discovered motifs, PWMs were generated in Python with the package logomaker (https://logomaker.readthedocs.io/en/latest/, version 0.8). Analyses and plots relative to Figures 5, S5 and S6H-I were generated in Python 3.9, with the exception of nucleotide contributions (DeepSHAP) which used Python 3.6. All other plots were generated using R or GraphPad Prism. Final plots were beautified on Adobe Illustrator.

### Data and code availability

Hight-throughput sequencing data will be made publicly available in GEO at the time of publication in a peer-reviewed journal. Code for the DaniO5P model is accessible at https://github.com/castillohair/DaniO5P.

### Supplemental Tables

**Table S1 - 5′ UTR MPRA library sequence information, relative mRNA 5′ UTR reporter abundances and sequencing statistics.** Sheet *Annotated_5*′ UTRs: Chromosome location, gene, transcript, consensus cluster and gene name information are given for each 5′ UTR entry. The full annotated 5′ UTR sequence is provided (merged_fasta_seq) as well as 5′ UTR length (utr_length). Promoter region genomic coordinates (consensus_cluster_start and consensus_cluster_end) and width (interquantile_width) are provided, as well as CAGEr derived expression values (tpm, tpm_dominant_ctss). Index denotes a unique identifier. Sheet *5*′ UTR_library: Chromosome location, gene, transcript, consensus cluster and gene name information are given for each 5′ UTR entry. Information on whether the 5′ UTR is a switching isoform (non-switching, maternal, zygotic, ZGA or postZGA) is included. The full annotated 5′ UTR sequence (merged_fasta_seq) and length (utr_length) is included. 5′ UTRs longer than 238 nt were split, the resulting sequences (split_utr_seq) and their length (utr_split_length) are also provided. Sheet *TPM_data_all_samples*: Identifier of 5′ UTR sequence assayed (insert_id) and relative transcript levels (TPM, transcript per million) for each sequence in each replicate sample. Sheet *Sequencing_stats*: Sequencing statistics for each high-throughput sequencing library are stated. Sheet *DNA_template_IVT_TPM*: Identifier of 5′ UTR sequence assayed (insert_id) and relative transcript levels (TPM, transcript per million) for the dsDNA PCR template used for generating the mRNA library by *in vitro* transcription (IVT); see Figure S1G. Sheet *Uninjected_IVT_RNA_lib_TPM*: Identifier of 5′ UTR sequence assayed (insert_id) and relative transcript levels (TPM, transcript per million) for the uninjected mRNA library generated by *in vitro* transcription (IVT); see Figure S1H.

**Table S2 – 5′ UTR MPRA RRS values.** Sheet *RRS_data_all_samples:* RRS values for each replicate measurement (repA, repB, repC) for all 5′ UTR sequence assayed (insert_id) at each time-point (2, 4, 6 and 10 hpf). Sheet *mean_log2_RRS_data:* mean of log_2_ transformed RRS values for each fraction at each developmental time-point. Information about the length of the 5′ UTR assayed (length_utr_assayed), the number of predicted upstream open reading frames (n_uORFS), the percent GC content (GC_content) and the predicted RNA structure (mxfold) is given. Sheet *Hierarchical clusters:* 5′ UTR sequences (insert_id) belonging to each of the nine hierarchical clusters (Cluster_ID) and respective mean log_2_ transformed RRS values for each fraction at each developmental time-point. See Figure 2 and STAR Methods.

**Table S3 – Data subsetting for motif enrichment analysis.** Sheet *Qs_RRS_HMW_10hpf:* information about the 5′ UTR sequences (insert_id) assayed, as well as mean log_2_ transformed RRS value for the HMW fraction at 10 hpf (mean_log2_RRS_HMW_10hpf), the nucleotide sequence of the 5′ UTR assayed (utr_seq_assayed), the 10-quantiles (Q) that each sequence belongs to based on RRS values, the length of the 5′ UTR sequence assayed (length_utr_assayed), the predicted number of upstream open reading frames (n_uORFs), the percent GC content (GC_content) and the predicted RNA structure (mxfold) is given. Sheet *TIS_nt_freqs:* nucleotide frequencies for positions −4 to −1 upstream of the sfGFP start codon for the bottom quantile (10%) (Q0_0.1) of the HMW at 10 hpf, the top quantile (10%) (Q0.9_1) of the HMW at 10 hpf and all assayed sequences in the MPRA. Sheet *Common_Upper_Q_80S:* 5′ UTR sequences commonly found in the upper quantile (top 10%) of the 80S fraction across the developmental time-point, grouped for motif-based sequence enrichment analysis. Sheet *Common_Lower_Q_80S:* 5′ UTR sequences in the lower quantile (bottom 10%) of the 80S fraction across the developmental time-course, grouped for motif-based sequence enrichment analysis. Sheet *Common_Upper_Q_LMW:* 5′ UTR sequences in the upper quantile (top 10%) of the LMW fraction across the developmental time-course, grouped for motif-based sequence enrichment analysis. Sheet *Common_Lower_Q_LMW:* 5′ UTR sequences in the lower quantile (bottom 10%) of the LMW fraction across the developmental time-course, grouped for motif-based sequence enrichment analysis. Sheet *Common_Upper_Q_HMW:* 5′ UTR sequences in the upper quantile (top 10%) of the HMW fraction across the developmental time-course, grouped for motif-based sequence enrichment analysis. Sheet *Common_Lower_Q_HMW:* 5′ UTR sequences in the lower quantile (bottom 10%) of the HMW fraction across the developmental time-course, grouped for motif-based sequence enrichment analysis. Sheet *Fuzzy_clusters_80S:* membership scores (ranging from 0 to 1) for each 5′ UTR sequence (insert_id) for each of the six fuzzy c-means clusters for the 80S fraction. Sheet *Fuzzy_clusters_LMW:* membership scores (ranging from 0 to 1) for each 5′ UTR sequence (insert_id) for each of the six fuzzy c-means clusters for the LMW fraction. Sheet *Fuzzy_clusters_HMW:* membership scores (ranging from 0 to 1) for each 5′ UTR sequence (insert_id) for each of the seven fuzzy c-means clusters for the HMW fraction. Sheet *Fuzzy_clusters_centroids: cluster* centroids derived from fuzzy c-means clustering. See also STAR Methods.

**Table S4 – List of motifs identified by motif enrichment analysis.** Sheet *Motif_enrichment_analysis:* consensus sequences of the motifs identified by the MEME motif-based sequence analysis tool (Bailey *et al*., 2015), associated E-values, information about which groups of 5′ UTR sequences they were enriched in (Sequence group) and know RNA-binding proteins that bind those motifs in other species (Motif ID and Motif Alt_ID). Sheet *Details_XSTREME:* additional information about motif-based search parameters. Sheet *PWMs_MEME_motifs*: position weight matrix of each motif outputted by XSTREME. Sheet *CNN_motifs*: names and consensus sequences of CNN model-discovered motifs. Sheet *PWMs_CNN_motifs*: position weight matrix of each motif outputted by the CNN model. Sheet *Tomtom_comparison*: p-, E- and q-values for the comparison of each CNN motif (Query) with each MEME motif (Target). Sheet *MAST_motif_matches*: MAST output file for search of MEME-derived motifs in 5′ UTR sequences of switching isoforms. Sheet *FIMO_motif_matches*: FIMO output file for search of MEME-derived motifs in 5′ UTR sequences of switching isoforms. Sheet *Length_model_parameters:* polynomial parameters of MRL length model. Sheet *Chrs_DaniO5P_model:* chromosomes and number of sequences in the training, validation, and test set of each model (n = 10). Sheet *Model_TPM_MRL_full_predictions*: MPRA sequences, along with the minimum total RNA TPM across replicates and time-points used as a data quality metric (min_TPM_input), calculated log2 MRL at all four time-points (columns starting with "log2_MRL”), calculated difference in total TPM (columns starting with "diff_log2_TPM”) at 4-2hpf, 6-2hpf, and 10-2hpf, length model predictions (columns starting with “pred_len”), ensemble CNN predictions (columns starting with "pred_cnn_ens”), and full DaniO5P predictions (columns starting with "pred_full”). See Figures 4, 5, S4, S5 and STAR Methods.

**Table S5 – Switching 5′ UTR analyses.** Sheet *Switching_5′ UTR _categories:* 5′ UTR entries (insert_id) for all switching 5′ UTR variants; top-ranked isoforms are indicated (category) as well as whether the isoform corresponds to a maternal, zygotic, ZGA or postZGA variant (type). Sheet *RSS_diff_m_z_HMW_10hpf:* difference (deltaRRS) in mean log_2_ RRS values for maternal and zygotic isoform pairs for the HMW fraction at 10 hpf; raking is based on deltaRRS values. Only 5′ UTR variants shorter than 239 nt were considered. Sheet *5′ UTRpairs_top_ranked_m:* mean log_2_ RRS values for top ranked maternal isoforms and their respective zygotic pairs (category); sequence of the 5′ UTR assayed and additional 5′ UTR features (uORF number, RNA folding score, GC content, length) are provided. Sheet *5′ UTRpairs_top_ranked_z:* mean log_2_ RRS values for top ranked zygotic isoforms and their respective maternal pairs (category). Sheet *RSS_diff_zga_postzga_HMW_10hpf:* difference (deltaRRS) in mean log_2_ RRS values for ZGA and postZGA isoform pairs for the HMW fraction at 10 hpf; raking is based on deltaRRS values. Only 5′ UTR variants shorter than 239 nt were considered. Sheet *5′ UTRpairs_top_ranked_zga:* mean log_2_ RRS values for top ranked ZGA isoforms and their respective postZGA pairs (category). Sheet *5′ UTRpairs_top_ranked_postzga:* mean log_2_ RRS values for top ranked postZGA isoforms and their respective ZGA pairs (category). See Figures 6, S6 and S7 and STAR Methods.

